# Quisinostat is a brain-penetrant radiosensitizer in glioblastoma

**DOI:** 10.1101/2022.11.09.515859

**Authors:** Costanza Lo Cascio, Tigran Margaryan, Ernesto Luna Melendez, James B. McNamara, Connor I. White, William Knight, Saisrinidhi Ganta, Zorana Opachich, Wonsuk Yoo, Nader Sanai, Artak Tovmasyan, Shwetal Mehta

**Affiliations:** Ivy Brain Tumor Center, Barrow Neurological Institute, Phoenix, Arizona, USA

## Abstract

In recent years, histone deacetylase inhibitors (HDACi) have garnered considerable interest for the treatment of adult and pediatric malignant brain tumors. However, owing to their broad-spectrum nature and inability to effectively penetrate the blood-brain barrier, HDACi have failed to provide significant clinical benefit to glioblastoma (GBM) patients to date. Moreover, global inhibition of HDACs results in widespread toxicity, highlighting the need for selective isoform targeting. While no isoform-specific HDACi are currently available, the second-generation hydroxamic acid-based HDACi quisinostat possesses sub-nanomolar specificity for class I HDAC isoforms, particularly HDAC1 and 2. Recently, we demonstrated that HDAC1 is the essential HDAC in GBM. Here, we provide the first report on the neuro-pharmacokinetic, pharmacodynamic and radiation-sensitizing properties of quisinostat in preclinical models of GBM. We demonstrate that quisinostat is a well-tolerated and brain-penetrant molecule that significantly extends survival when administered in combination with radiation *in vivo*. The pharmacokinetic-pharmacodynamic-efficacy relationship was established by correlating free drug concentrations and evidence of target modulation in the brain with survival benefit. Together, these data provide a strong rationale for clinical development of quisinostat as a radiosensitizer for the treatment of GBM.

## INTRODUCTION

HDAC inhibitors (HDACi) are a successful example of epigenetic therapy, with five inhibitors currently FDA-approved for the treatment of different hematological malignancies, and a growing number of agents currently in different stages of clinical testing for a variety of cancers (1). Over the past 15 years, numerous HDACi have been studied preclinically in the realm of neuro-oncology, three of which (vorinostat, romidepsin and panobinostat) have been tested in clinical trials for patients with primary and recurrent glioblastoma (GBM) (2). Unfortunately, all HDACi have failed to significantly prolong survival in this population of patients to date. The disappointing clinical results with HDACi in the treatment of GBM can be attributed to inadequate disease modeling at the preclinical level, poor blood-brain barrier penetration, and limited central nervous system (CNS) pharmacokinetic profiling (2-4).

HDACi that have been tested for GBM in the clinic so far have been broad-spectrum (pan-HDACi), which non-selectively target multiple human HDAC isoforms (2). Considering that HDACs retain essential functions for cell homeostasis across different tissues, pan-HDACi are associated with serious adverse events – a notable example being the hydroxamic acid panobinostat (5-7). Thus, HDACi-induced toxicities significantly restrict the therapeutic window of these drugs for the treatment of CNS malignancies. However, it has been suggested that improved drug target selectivity typically leads to a superior safety profile, and this may hold true for HDACi as well (8, 9).

Isoform selectivity of HDACi is an important consideration given that not all HDAC enzymes are equally expressed in GBM, and that the specific roles of individual HDAC isoforms in these tumors are not well understood (10). We recently uncovered the functional importance of HDAC1 in GBM, an HDAC isoform whose expression increases with brain tumor grade and is correlated with decreased survival (11). We found that HDAC1 function is essential for the survival of glioma stem cells (GSCs) and that its loss is not compensated for by its paralogue HDAC2 or other HDACs. Importantly, we demonstrated that loss of HDAC1 alone significantly prolonged survival *in vivo* – providing a rationale for the development of isoform-selective HDACi for the treatment of GBM (11).

While no HDAC1-selective agents are currently available, quisinostat (JNJ-26481585) is a second-generation HDACi that is highly selective towards class I HDACs and harbors marked potency towards HDAC1 (IC_50_: 0.1 nM) (8). Quisinostat has been shown to exhibit potent antitumor activity in preclinical models of different cancers and has been studied in phase I/II clinical trials for ovarian and hematological malignancies (12, 13). However, there have been inconclusive reports on the survival benefits of quisinostat treatment in orthotopic settings for brain tumors (GL261, SHH medulloblastoma, DIPG) (14-16). It is worth nothing that these studies lacked direct pharmacokinetic and pharmacodynamic data and/or were conducted in flank tumor models.

Here, we report detailed CNS and brain tumor pharmacokinetic (PK), pharmacodynamic (PD), and radiation-sensitizing properties of quisinostat in preclinical models of human GBM. We find that quisinostat inhibits the growth of multiple GSC lines, induces histone hyperacetylation, DNA damage, cell death, and cell cycle arrest. Importantly, we demonstrate that quisinostat is a brain-penetrant molecule that can significantly extend survival of an orthotopic patient-derived xenograft model of GBM when combined with radiation therapy. Together, our results reveal that quisinostat is a potent radiosensitizer, providing a rationale for clinical development of quisinostat in combination with radiation for GBM treatment.

## RESULTS

### Quisinostat is cytotoxic and induces stable global hyperacetylation in patient-derived GSCs

Quisinostat (JNJ-26481585) is a second-generation hydroxamic acid that harbors remarkable selectivity towards class I HDACs with a biochemical half maximal inhibitory concentration (IC_50_) of 0.1 nM for HDAC1 (13). Quisinostat has demonstrated *in vitro* efficacy across multiple human cell lines derived from aggressive pediatric brain tumors (diffuse intrinsic pontine glioma, sonic hedgehog medulloblastoma) (15, 16) but not GBM cell lines. To determine the cytotoxic effects of quisinostat, we performed a dose-titration cell viability assay in 7 patient-derived GSC lines (BT145, GB187, GB239, GB282, GB71, GB82, GB126) and one long-term serum-grown human GBM cell line (U87). As GBMs display a high degree of intra-tumoral heterogeneity, we used GSCs derived from both primary and recurrent GBMs that harbored distinct genetic mutations or aberrations, growth rates, *MGMT* promoter methylation status and gene expression profiles (Supplemental Figure 1). Cell lines were treated with various concentrations of quisinostat (10-1000 nM), and cell viability was measured 3-5 days later (Figure 1 A-B). The cellular IC_50_ for quisinostat in all lines was in the low nanomolar range (50 – 100 nM), demonstrating greater potency (< 1 µM) compared to other pan-HDACi (VPA, TSA, vorinostat, entinostat) that have been tested on GBM cells in previous studies (17-22). Treatment with quisinostat at IC_50_ concentrations induced significant inhibition of proliferation (Ki67) and an increase in programmed cell death (cleaved caspase 3) in two different GSC lines (Figure 1 C-D). These results indicate a cytotoxic effect of quisinostat on GSC cultures.

**Figure 1.**
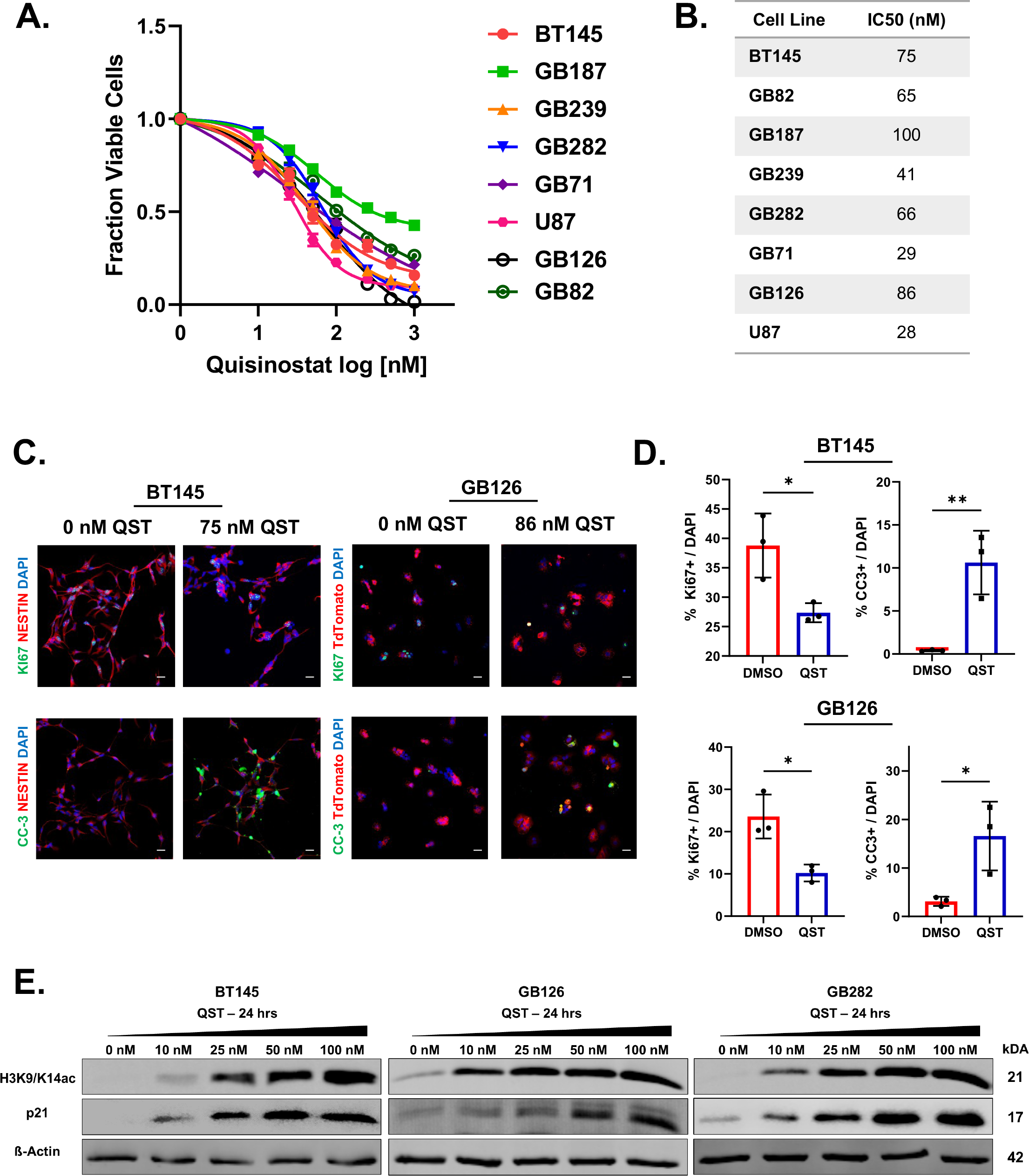
Quisinostat exhibits low nanomolar efficacy against human glioma stem cell cultures. (A) Dose-response curves with quisinostat (10-1000 nM). Cell viability was measured across 7 patient-derived GSCs and one serum-grown long-term glioma line (U87) 3-5 days after treatment with quisinostat. (B) Table illustrating the half maximal inhibitory concentration (IC_50_) of quisinostat for each cell line tested. (C) Immunofluorescence staining of GSC lines BT145 (left) and GB126 (right) 72 hours after treatment with quisinostat at the IC_50_ concentrations. Control and drug-treated cells were stained for Ki67 and cleaved caspase-3 to assess cell proliferation and cell death respectively. (D) Quantification of Ki67-positive and cleaved caspase-3 positive cells in BT145 (top) and GB126 (bottom) 72 hours after treatment with quisinostat (n=3). (E) Representative immunoblots showing dose-dependent increase in histone H3K9/14 acetylation and p21 levels in GSC lines BT145, GB126 and GB282 after 24-hour treatment with quisinostat. QST = quisinostat. Error bars indicate SEM. * *p* < 0.05, ** *p* < 0.01 . Magnification, 20x; scale bars, 20 μM. *P* values were determined using the unpaired 2-tailed t-test.

We next investigated the cellular effects of quisinostat on histone acetylation dynamics in GSCs. Immunoblot analysis of lysates from three independent GSC lines (BT145, GB126, GB282) treated with increasing concentrations of quisinostat (range: 10-100 nM) revealed a significant dose-dependent increase in histone H3 acetylation at lysines 9 and 14 (H3K9/14ac), indicative of target engagement given that histone acetylation is primarily regulated by HDAC1 and HDAC2 (Figure 1E) (23). In agreement with the results shown in Figure 1C-D, we observed a dose-dependent increase in expression of p21 – tumor suppressor protein and a key negative regulator of the cell cycle (Figure 1E). These results suggest that quisinostat induces global changes histone hyperacetylation, increasing chromatin accessibility, and promotes cell cycle arrest in GSCs.

### Quisinostat induces sustained levels of DNA damage

Previous studies have demonstrated that several HDACi can act as DNA-damaging agents in malignant cells (24-26). To test DNA damaging effects of quisinostat in GBM cells, we treated two independent GSC lines with quisinostat for 72 hours at their IC_50_ concentrations and analyzed changes in levels of phosphorylated histone H2AX (γH2AX), a marker for DNA double-strand breaks (DSB) (27). We found that there were significantly more γH2AX foci per cell in both cell lines post-treatment compared to DMSO-treated controls (Figure 2A-B).

**Figure 2.**
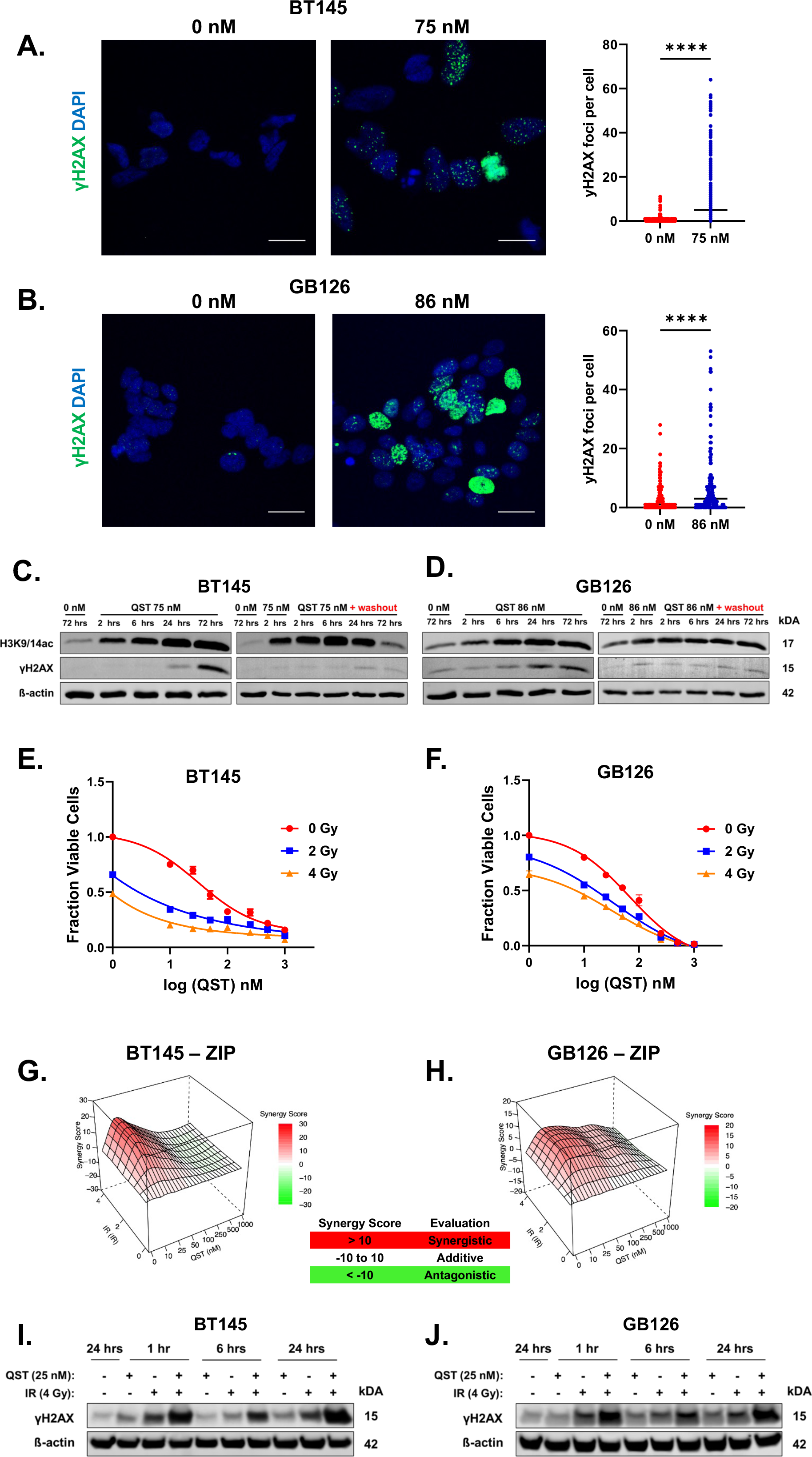
Quisinostat sensitizes GSCs to ionizing radiation. (A-B) Immunofluorescence staining of BT145 (A) and GB126 (B) showing an increase in γH2AX foci 72 hours after treatment with quisinostat but not DMSO-treated cells. Average number of γH2AX quantified in each cell per treatment condition are shown to the right for BT145 and GB126. (C-D) Representative immunoblots demonstrating that quisinostat treatment in (C) BT145 and (D) GB126 results in accumulation of γH2AX over time (left) and but does not after drug washout (right). (E-F) Dose response curves combining quisinostat and ionizing radiation treatment in (E) BT145 (F) and GB126. (G-H) Matrices illustrating the ZIP synergy scores when combining quisinostat with increasing doses of radiation in (G) BT145 and (H) GB126. (I-J) Representative immunoblots showing protein levels of γH2AX in (I) BT145 and (J) GB126 1, 6 and 24 hours after treatment with either quisinostat alone (25 nM), radiation alone (4 Gy) or both quisinostat and radiation (25 nM and 4 Gy). QST = quisinostat, IR = ionizing radiation. For each cell line, the data are compiled from at least three independent experiments. Magnification, 63x; scale bars, 20 μM.

We next sought to understand the temporal dynamics of quisinostat-induced DNA DSB in GSCs. To do this, we treated two different GSC lines at their respective IC_50_ values for quisinostat (75 nM for BT145 and 86 nM for GB126) and utilized immunoblotting to observe changes in the levels of γH2AX at 2, 6, 24 and 72 hours after treatment with the drug (left panels, Figures 2C-D). We observed that histone H3 hyperacetylation increased over time after continuous exposure to quisinostat, while γH2AX accumulated more gradually with levels peaking at 72 hours after treatment in both cell lines (left panels, Figures 2C-D). To characterize the reversibility of the expected PD effect, we performed cell washout experiments at different timepoints after drug removal (right panels, Figures 2C-D). To determine whether histone H3 hyperacetylation and DNA damage persisted after acute exposure to quisinostat, we replaced drug-spiked media with drug-free media 2 hours post-treatment (right panels, Figures 2 C-D). Washout experiments demonstrated that quisinostat target engagement (assessed through changes in histone H3K9/14 acetylation) persist after its removal from the media. Although histone H3 acetylation levels in BT145 were still higher at 24 hours relative to DMSO-treated samples, they dramatically decreased 72 hours after drug washout (right panel, Figure 2C).

Unlike what we observed in BT145, in GB126 upon drug washout histone H3 acetylation did not decrease over time and remained relatively stable in the absence of the drug in the media (right panel, Figure 2D). Moreover, in both cell lines, we found that γH2AX levels did not dramatically increase in timepoints following drug washout (right panels, Figures 2C-D). Hence, these results demonstrate that quisinostat-induced inhibition of class I HDAC activity is relatively stable (∼24-72 hrs) and capable of inducing a prolonged PD effect (histone acetylation) in GSCs, even after short-term incubation with the drug. The lack of significant accumulation in DNA DSB suggests that continuous or persistent drug exposure is necessary for quisinostat to induce substantial DNA damage in GSCs (right panels, Figure 2 C-D). To the best of our knowledge, this is the first report that indicates that quisinostat can act as a potent DNA-damaging agent in cancer cells *in vitro*.

### Kinetics of intracellular uptake of quisinostat in GSCs

To further understand the kinetics of drug-target engagement in GSCs, we harvested media and BT145 cells at 0, 2, 6, 10 and 24 hour timepoints with or without removal of quisinostat (75 nM). At each timepoint, we measured the intracellular levels of the drug as well as its concentrations in the cell media. With continuous drug exposure (no washout), the intracellular levels of quisinostat in BT145 gradually increased throughout the incubation period, reaching equilibrium by 10 hours (blue line, Supplementary Figure 2A). However, the levels of quisinostat decreased in cell media over time (orange line with triangles, Supplementary Figure 2A), suggesting that the drug may not be very stable in GSC culture media relative to baseline drug levels (orange line with circles, Supplementary Figure 2A). By contrast, in washout experiments (performed 2 hours after spiking cell media with the drug), minimal quisinostat levels were measured intracellularly after removal of the drug at all time points (blue line, Supplementary Figure 2B). Hence, our results indicate that the low intracellular levels of drug after washout (<10 nM; blue line, Supplementary Figure 2B) are sufficient to inhibit HDAC activity, albeit short-lived compared to continuous drug exposure (Figure 2C).

### Quisinostat treatment sensitizes GSCs to ionizing radiation in vitro

In addition to their use as single-modality anticancer agents, there is preclinical evidence in other cancers that HDACi may be effective in enhancing radiosensitivity of tumor cells when combined with radiation therapy (24, 28-31). We hypothesized that the accumulation of DNA damage induced by quisinostat in combination with radiation treatment may synergistically reduce GSC viability. To examine this, we treated two cell lines (BT145 and GB126) with increasing nanomolar doses of quisinostat (10-1000 nM) and increasing doses of IR (2-4 Gy) (Figure 2 E-F). Across both cell lines, combination treatment resulted in greater cytotoxicity compared to independent treatment with quisinostat or radiation (Figure 2E-F). We then analyzed our combinatorial dose-response cell viability data using SynergyFinder, an application that assigns synergy scores using various major reference models (32). The zero interaction potency (ZIP), BLISS and Loewe model synergy matrices indicated that the greatest synergy was attained when combining IR with the lowest doses of quisinostat tested (10-25 nM) (Figure 2 G-H, Supplemental Figure 3A-B). Immunoblotting analysis of BT145 and GB126 cells treated with 25 nM quisinostat and/or 4 Gy radiation revealed that combinatorial treatment resulted in significantly higher levels of γH2AX compared to either treatment alone (Figure 2 I-J). Together, these data demonstrate that low nanomolar doses of quisinostat can enhance sensitivity of GSCs to radiation treatment.

### Determination of optimal route of administration for quisinostat in vivo

Considering that the efficacy of quisinostat in preclinical models of brain tumors remains controversial and that previous studies employed various drug delivery methods, we sought to understand how different routes of administration impact the bioavailability of quisinostat in mice. Three routes of administration were compared to determine the optimal dosing route to obtain highest plasma exposure to quisinostat over time. We treated athymic nude mice with a single dose of quisinostat (10 mg/kg) through either intraperitoneal (IP), subcutaneous (SQ) or oral delivery (OG) and collected blood at 0.5, 1, 2, 4, 6, 8 and 24 hours post-dosing from individual mice for PK analysis (Figures 3A). LC/MS analysis revealed that regardless of administration route, quisinostat was systemically cleared within 24 hours of dosing (Figure 3B). IP and SQ injections of quisinostat resulted in higher plasma exposure over time compared to OG delivery (AUClast 1202.2, 1187.1 and 106.0 respectively; Figure 3B). Therefore, IP delivery was established to be the optimal route of administration to maximize the bioavailability of quisinostat in athymic nude mice.

**Figure 3.**
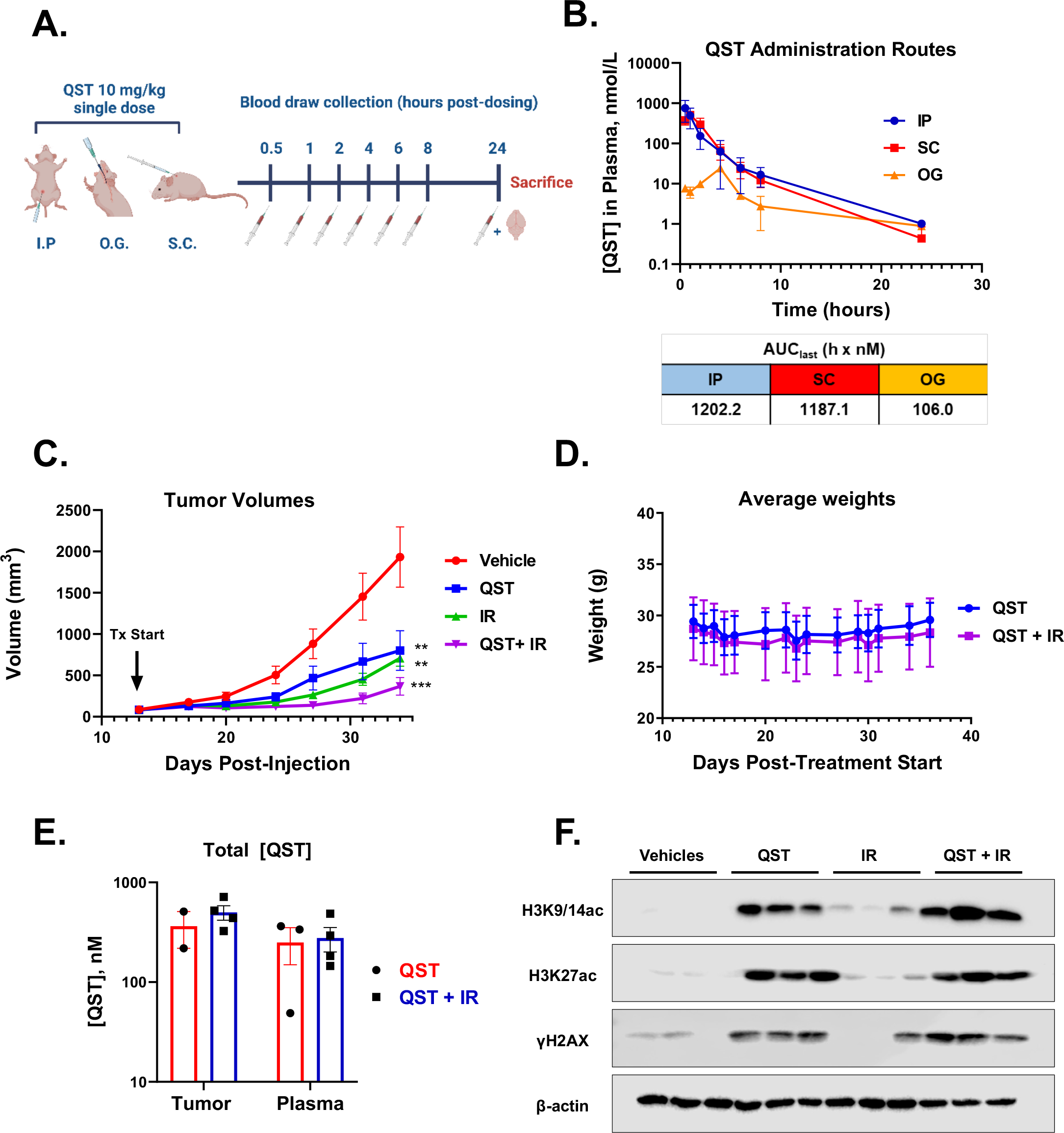
Quisinostat is effective in slowing tumor growth in a flank model of human GBM. (A) Schematic illustrating experimental design Athymic nude mice were treated with a single dose of quisinostat (10 mg/kg) through either intraperitoneal (I.P.), subcutaneous (S.C.) or oral gavage (O.G.). Blood samples were collected at 0.5, 1, 2, 4, 6, 8 and 24 hours post-dosing and analyzed by LC-MS/MS. (B) Total plasma concentration-time curved for quisinostat administered through various routes. Area under the curve values (AUC, h×nM,) were calculated for each route to illustrate plasma quisinostat exposure (bottom). (C) Weekly volume measurements of U87 flank tumors from mice treated with vehicle, 10 mg/kg quisinostat, radiation alone (6 Gy) or combination treatment (6 Gy and 10 mg/kg quisinostat) (n=10 in each cohort). (D) Average weights of mice treated with quisinostat (dosed at 10 mg/kg) (blue) or combination therapy (purple) throughout the entire duration of the study. (E) Total levels of quisinostat in plasma and flank tumors of quisinostat- and combination-treated mice (n=2-4 per cohort). (F) Immunoblotting of protein lysates derived homogenized flank tumors from each cohort (n=3 per group). Membranes were probed for H3K9/14ac, H3K27ac, γH2AX and B-actin. QST = quisinostat, IR = ionizng radiation. Error bars indicate SEM. * p < 0.05, ** p < 0.01, *** p < 0.001 n.s., P values were calculated the ordinary one-away ANOVA with Dunnett’s multiple comparisons test.

### Quisinostat inhibits tumor growth in a flank model of human GBM

We next examined if quisinostat treatment alone or in combination with IR would be effective in slowing tumor growth in a flank model of GBM (U87). Tumor-bearing mice receiving radiation were treated with 2 Gy fractions on MWF 2 hours after dosing with either vehicle solution or quisinostat (10 mg/kg), for a total dose of 6 Gy. Following completion of radiation treatment, mice continued to receive vehicle or quisinostat until the tumors reached the maximum volume threshold (Figure 3C). Quisinostat monotherapy significantly reduced tumor volume compared to vehicle-treated mice (Figure 3C). Combination treatment was more effective in reducing tumor growth than either quisinostat or radiation therapy alone, the average tumor volume being ∼ 4.5 fold smaller compared to tumors from vehicle-treated mice at the end of the study (Figure 3C). We found that quisinostat, even in combination with radiation, was well-tolerated throughout the entirety of the treatment study when dosed at 10 mg/kg intraperitoneally MWF, with no significant loss in average weights throughout the entire regimen (Figure 3D).

PK analyses of quisinostat- and combination-treated mice demonstrated that average total quisinostat concentrations were higher in the flank tumors (∼433 nM) than in the plasma samples (∼300 nM) (Figure 3E). There was no significant difference in the total levels of quisinostat between monotherapy and combination cohorts (Figure 3E). Immunoblotting confirmed significant increase in histone H3 acetylation at lysines 9, 14, and 27 in mice treated with quisinostat alone or in combination with radiation (H3K9/14ac, H3K27ac; Figure 3F). Moreover, we observed that all quisinostat treated tumors expressed high levels of γH2AX – indicative of the presence of double stranded DNA breaks – relative to vehicle-treated controls (Figure 3F). These results suggest that 10 mg/kg dosing of quisinostat is effective in reducing tumor burden, induces the intended PD effects in glioma cells and corroborate our previous *in vitro* findings that quisinostat acts as a potent DNA-damaging agent.

### Inter-species differences in the stability of quisinostat

Hydroxamic acid-based compounds have been reported to display poor stability and high plasma clearance due to the presence of arylesterases and carboxylesterases in rodent blood (33). Therefore, we measured the stability of quisinostat in mouse plasma and brain to determine whether drug degradation occurred during our sample preparations and equilibrium dialyses for the determination of unbound drug level (performed at 37 °C). We found that quisinostat exhibited significant instability in mouse plasma during an 8-hour incubation time with a half-life of ∼1 hour (Figure 4A). Quisinostat degradation was also observed in mouse brain homogenate (green line, Figure 4B). However, perfusion of mice prior to collection of their brains prevented rapid drug degradation in the brain matrix (blue line, Figure 4B). This indicates that quisinostat instability in the brain is most likely related to the enzymes that are present in the mouse plasma. Degradation of quisinostat can also be significantly constrained if plasma and brain samples are stored in a refrigerator at 4 °C (Supplementary Figure 4A). All subsequent sample preparations were therefore performed on ice-cold baths to avoid degradation of quisinostat in our PK analyses. Importantly, we demonstrate that the stability of quisinostat can also be prolonged in the presence of the carboxylesterase inhibitor bis(p-nitrophenyl) phosphate (BNPP, Supplementary Figure 3B). Hence, the dialyses of mouse plasma and brain samples were executed in the presence of BNPP to maximize quisinostat stability at 37°C. Interestingly, the drug was completely stable in human plasma and brain homogenate at 37°C (Figure 4C-D). The observed stability is probably due to the absence of esterases in human matrices, which are responsible for degradation of quisinostat (33). Since we employed quisinostat in several *in vitro* studies (Figures 1-2), the stability of the molecule was tested also in the cell media used to culture GSCs (neurobasal medium, refer to Methods). We demonstrate that ∼70% of quisinostat stays intact in the cell media over a 24-hour incubation period at 37 °C (Supplemental Figure 4C). These results are in line with the data obtained in the intracellular uptake experiments illustrated in Supplemental Figure 2A.

**Figure 4.**
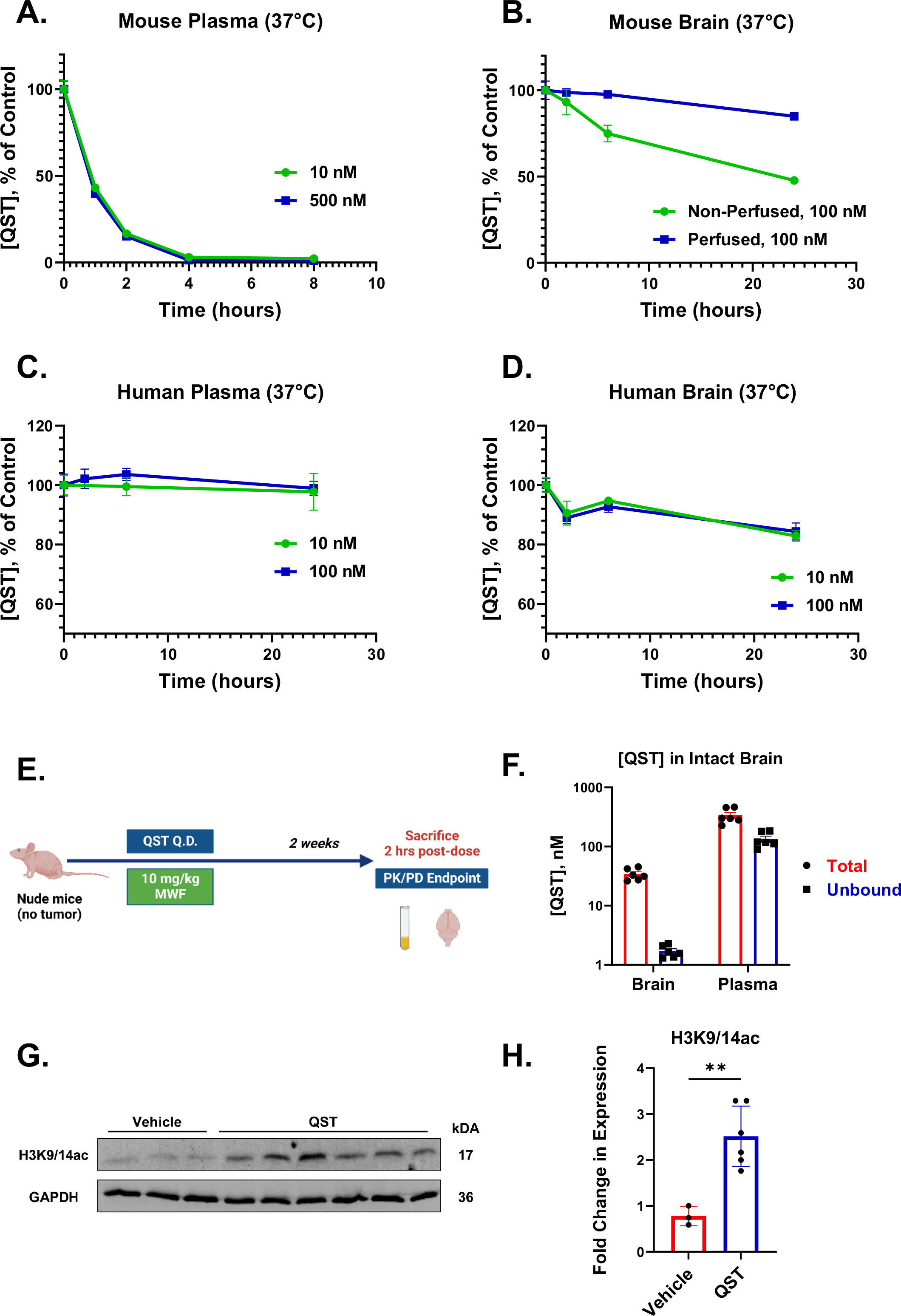
Quisinostat is a brain-penetrant HDACi. (A) Stability of quisinostat (10 nM and 500 nM) in mouse plasma. (B) Stability of quisinostat in mouse non-perfused and perfused brain homogeneate (1/7 w/v in PBS). (C) Stability of quisinostat in human plasma. (D) Stability of quisinostat in human brain homogeneate. (E) Schematic illustrating the design of the treatment study in non-tumor bearing athymic nude mice. (F) Total (red) and unbound (blue) levels of quisinostat in normal brain tissue in quisinostat-treated mice (n=6 per cohort). (G) Immunoblotting of protein lysates derived homogenized brains from each cohort (n=3 for vehicle cohort, n=6 for quisinostat cohort). Membranes were probed for H3K9/14ac and GAPDH. (H) Quantification of expression of H3K9/14ac (normalized to GAPDH) in quisinostat and vehicle-treated homogenized brain samples. QST = quisinostat. ** p < 0.01. P values were determined using the unpaired 2-tailed t-test. Values are the mean of triplicate measurements and error bars indicate SEM. QST = quisinostat.

### Pharmacokinetics and pharmacodynamics of quisinostat in the normal central nervous system

Insufficient drug exposure in the brain is one of the major hurdles in the treatment of brain tumors. Hence, we determined the PK profile of quisinostat in the normal central nervous system (CNS) by treating athymic nude mice with 10 mg/kg quisinostat on a MWF schedule for two weeks. On the last day of treatment, mice were sacrificed 2 hours after dosing with quisinostat and blood and intact brains were harvested (Figure 4E). Each brain hemisphere was processed separately to perform matched PK and PD analyses from the same animal. We found that although the average unbound levels of quisinostat in the brain were >70-fold lower than in the plasma, the pharmacologically active unbound drug concentration in the brain (∼ 1.7 nM) was over 15 times higher the biochemical IC_50_ for HDAC1 (0.1 nM) (Figure 4F). For PD analyses, we homogenized entire hemispheres to obtain whole tissue protein lysates from each mouse and assessed changes in histone H3 acetylation levels using immunoblotting. We confirmed that relative to vehicle-treated animals, the levels of H3K9/14 acetylation were significantly increased in the brain tissue of quisinostat-treated mice (n=6) (Figure 4G-H). Our results therefore reveal that quisinostat is a brain-penetrant HDACi that exhibits clear on-target PD activity in normal CNS cells. We also established a direct correlation between PK and PD modulation *in vivo*, demonstrating that the free unbound levels of quisinostat in the brain can induce substantial histone H3 hyperacetylation (Figure 4G-H).

### Quisinostat is a potent radio-sensitizer in an orthotopic patient-derived xenograft model of GBM

To establish the PK-PD correlation and efficacy in an orthotopic PDX model of GBM, we implanted GB126 cells in the brains of athymic nude mice and began treatment once the tumors started growing exponentially. Tumor-bearing mice were treated with quisinostat (10 mg/kg), with or without radiation, on a MWF schedule (Figure 5A). As described previously, IR was delivered locally to the brain in 2 Gy fractions 2 hours after being dosed with vehicle solution or quisinostat, for a cumulative delivery of 6 Gy. To assess acute PD effects and drug levels after short-term quisinostat treatment, 10 mice from each experimental cohort were sacrificed 3 hours after receiving the third dose of vehicle solution or quisinostat in the first week of treatment (Figure 5A). Plasma, tumors, and brain tissue contralateral to the tumors were collected from each mouse and subsequently processed for PK and PD analyses. As shown in Figures 5B and 5C, total and unbound levels of quisinostat accumulated in tumor tissue compared to contralateral brain tissue. However, unbound quisinostat levels in the tumors were up to 600-fold above the biochemical IC_50_ for HDAC1 inhibition. Importantly, although quisinostat levels in contralateral brain tissue were significantly lower than those measured in the tumors, they were 12-fold higher the biochemical IC_50_ of HDAC1, which ensures target inhibition in infiltrating tumor cells distant from the tumor core. No significant differences in drug accumulation were observed between the monotherapy and combination therapy cohorts. To assess PD modulation, we homogenized brain tumors to obtain whole tissue protein lysates from each mouse and quantified changes histone H3 acetylation levels using immunoblotting. We confirmed that relative to vehicle-treated animals, the levels of H3K9/14 and H3K27 acetylation increased in monotherapy- and combination-treated mice relative to vehicle-treated mice (Figure 5D). Moreover, we detected elevated levels of γH2AX in combination-treated tumors, corroborating our previous findings that combining quisinostat and radiation results in high levels of DNA damage in GSCs *in vitro* (Figure 2I-J).

**Figure 5.**
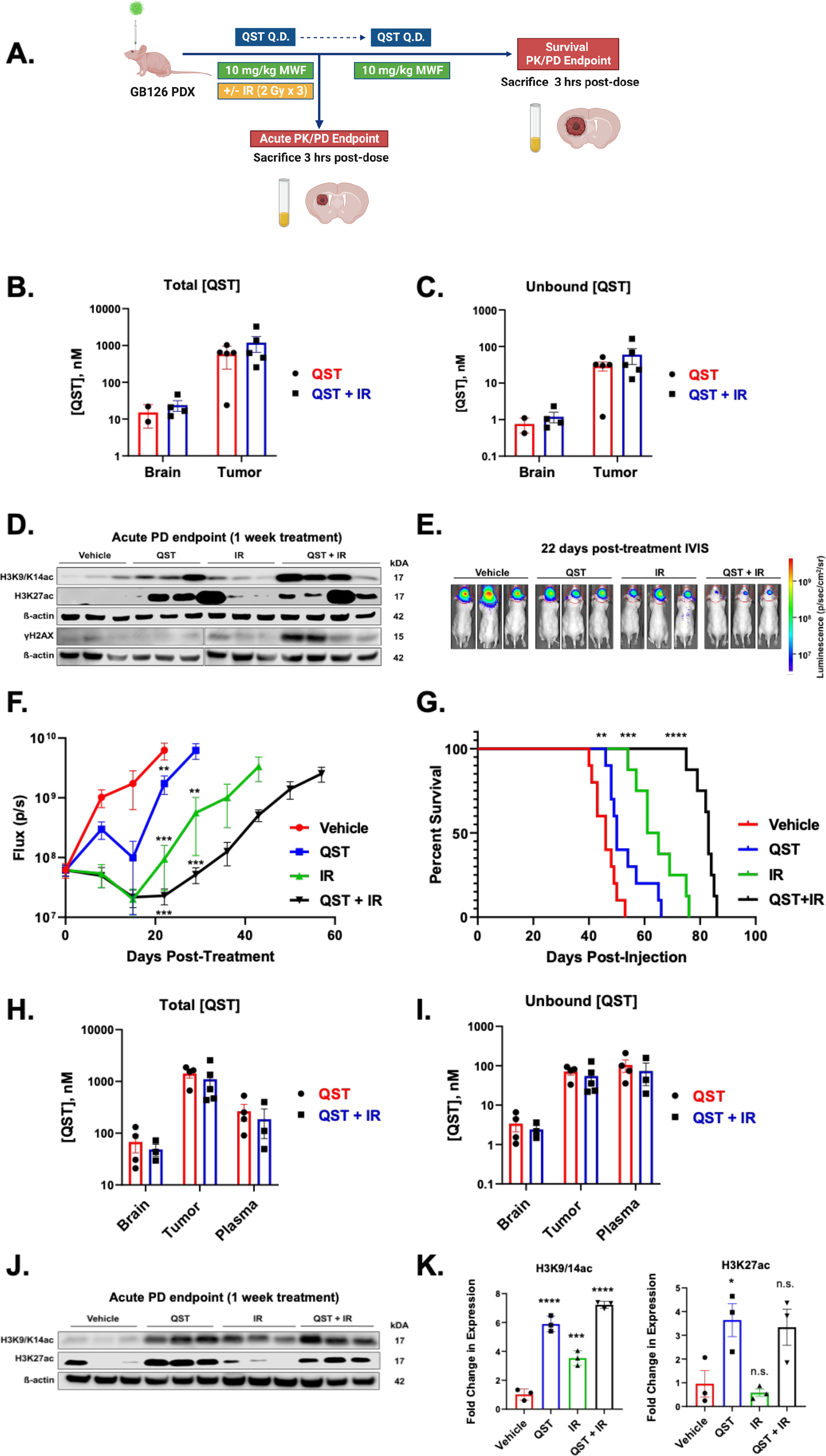
Quisinostat is as a potent radiosensitizer in an orthotopic patient-derived xenograft model of GBM. (A) Schematic illustrating the experimental design of the treatment study in an orthotopic PDX model of GBM (GB126). (B-C) Total (B) and unbound (C) levels of quisinostat in tumor tissue and brain tissue contralateral to the tumor in quisinostat- and combination-treated mice (n=2-5 per cohort). (D) Immunoblotting of protein lysates derived homogenized brain tumors from each cohort (n=3-4 per cohort). Membranes were probed for H3K9/14ac, H3K27ac, γH2AX and B-actin. (E) Representative images of representative heatmap of bioluminescence intensity across all treatment cohorts 22 days post-treatment initation. (F) Average photon flux (p/s) measured through bioluminescence imaging across all cohorts throughout the entire duration of the treatment. (G) Kaplan-Meier survival analysis of vehicle-, quisinostat- (10 mg/kg), radiation- (6 Gy) or combination-treated (6 Gy total radiation and 10 mg/kg quisinostat) mice. (H-I) Total (H) and unbound (I) levels of quisinostat in tumor tissue and brain tissue contralateral to the tumor in quisinostat- and combination-treated mice (n=4-5 per cohort). (J) Immunoblotting of protein lysates derived homogenized brain tumors from each cohort (n=3 per cohort). Membranes were probed for H3K9/14ac, H3K27ac and B- actin. (K) Normalized levels of H3K9/14ac and H3K27ac protein in all cohorts are shown to the right. QST = quisinostat. Error bars indicate SEM. * p < 0.05, ** p < 0.01, *** p < 0.001, **** p < 0.0001 n.s., not significant. P values were calculated using unpaired 2-tailed t-test and Kaplan-Meier method with the Mantel-Cox log-rank test.

We additionally questioned whether quisinostat, either alone or in combination with radiation therapy, could extend survival of tumor-bearing mice. Following completion of the radiation regimen, mice continued to receive quisinostat at 10 mg/kg or vehicle solution on MWF until the end of the study, as determined by large tumor burden and onset of neurological symptoms (Figure 5A). Bioluminescence imaging demonstrated that quisinostat monotherapy or its combination with radiation significantly reduced tumor burden compared to vehicle or radiation-only cohorts (Figure 5E-F). While quisinostat monotherapy slowed tumor growth, it only resulted in an average increase in survival of 4 days (*p* < 0.01) relative to vehicle-treated mice, and radiation treatment alone increased median survival by an average of 17 days (*p* < 0.001, Figure 5G). However, combining quisinostat with radiation treatment led to a substantial increase in median survival (37 days, *p* < 0.0001) compared to vehicle and quisinostat or radiation-monotherapy cohorts (Figure 5G). These data suggest that while quisinostat treatment alone produces a modest therapeutic benefit, combinatorial treatment with fractioned doses of radiation unveils that quisinostat acts a potent radiosensitizer that significantly prolongs survival in an orthotopic PDX model of human GBM (Figure 5G).

All the mice in each cohort utilized in the survival study were utilized for end-point PK and PD analyses once moribund. As described above, plasma, tumor and contralateral brain tissues were harvested 3 hours after dosing with 10 mg/kg quisinostat, allowing for a direct comparison of long-term treatment with the PK/PD data collected from acute (1 week) treatment with quisinostat or combination therapy (Figure 5B-D). As shown in Figure 5H-I, PK analyses demonstrated that unbound quisinostat accumulated in the tumors (average ∼71.4 nM) and peritumoral brain tissue (average ∼3.4 nM) over time. There were no significant differences in total or unbound drug concentrations in tumor or brain tissue between the monotherapy or combination therapy cohorts (Figure 5 H-I). Immunoblot analysis of resected tumor samples confirmed that quisinostat induced substantial histone H3 hyperacetylation in brain tumors compared to untreated animals, consistent with an *in vivo* on-target effect (Figure 5J-K). Our results establish that quisinostat is a brain-penetrant drug that accumulates in both normal brain and tumor tissue.

### Combined quisinostat and radiation treatment induces neuronal differentiation and cell cycle arrest

To better understand the downstream effects of the various treatment regimens, we performed RNA sequencing (RNA-seq) to analyze the transcriptomes of GB126 tumors of mice treated with either acute (1 week) or prolonged quisinostat monotherapy (QST), IR or combination treatment (QST+IR). Acute quisinostat (1 week) treatment resulted in significant transcriptional changes relative to vehicle controls (908 upregulated and 396 downregulated genes) (Supplemental Figures 5A-C). Conversely, long-term quisinostat monotherapy led to modest changes in gene expression, (341 upregulated and 147 downregulated genes) (Figures 6A-B). Short-term fractionated radiation therapy (2 Gy x 3 MWF, Figure 5A) resulted in minimal changes in gene expression compared to vehicle treatment (Figures 6A-B). Conversely, long-term combination treatment induced much more pronounced changes in gene expression with 1,208 upregulated and 465 downregulated genes (FDR < 0.05, 2-fold, Figures 6A-B). These data suggest that while positive and negative changes in the expression of some genes are shared across all cohorts in end-stage tumors (Figure 6A), the transcriptomes of the combination-treated tumors are largely distinct from tumors that received either drug or radiation treatment alone (Figure 6C).

**Figure 6.**
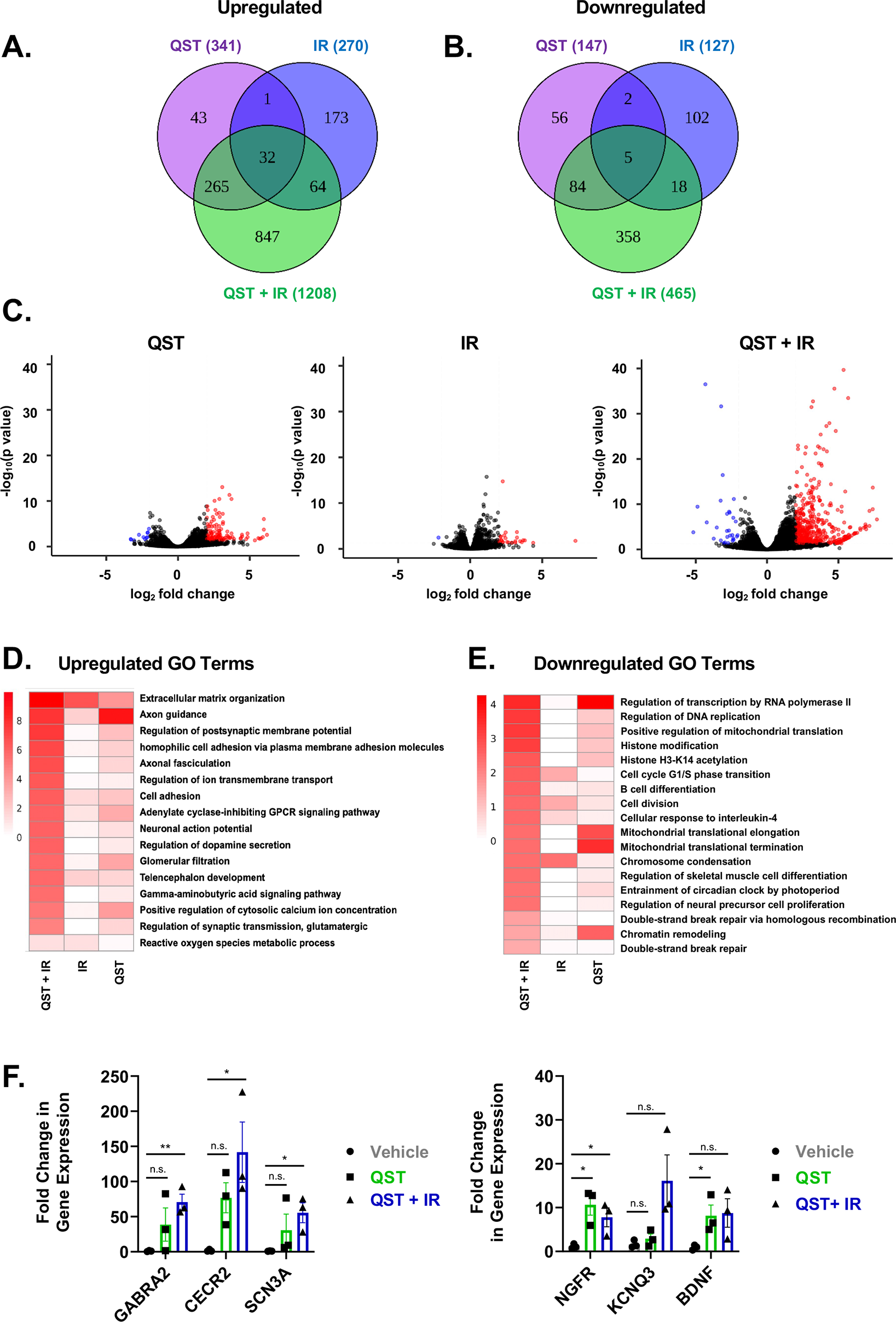
Combined treatment or quisinostat and radiation induces cell cycle arrest neuronal differentiation *in vivo*. (A-B) Venn diagrams showing the overlap in genes either upregulated (A) or downregulated (B) in response to quisinostat (QST) monotherapy, radiation treatment (IR) or combination treatment (QST + IR). Gene numbers in each section are shown in parentheses. (C) Volcano plots showing the –log10 (p value) and log2 fold change for transcripts detected by RNA-seq analysis of end-point orthotopic GB126 xenograft tumors treated with QST (left), IR (center), or QST + IR (right). Significantly up- and downregulated genes (FDR < 0.05, 2-fold) are marked in red and blue, respectively. (D-E) Gene ontology analysis of genes upregulated (D) or downregulated (E) in GB126 tumors due to QST, IR or QST + IR treatment. (F) RT-qPCR analysis of the expression of neuronal markers in GB126 tumors treated with either QST monotherapy or combination therapy. QST = quisinostat. Error bars indicate SEM. * p < 0.05, ** p < 0.01, n.s., not significant P values were calculated using unpaired 2-tailed t-test.

To determine the biological processes that are differentially enriched in treatment groups, we performed gene ontology (GO) analysis on both upregulated and downregulated datasets from each treatment group (34). In acutely treated tumors, GO analysis confirmed that combination treatment upregulated genes related to responses to cellular stress (oxidative stress, wound healing, unfolded protein response), apoptosis and cellular differentiation (Supplemental Figure 5D). Quisinostat monotherapy resulted in similar responses, albeit not as pronounced (Supplemental Figure 5D). Furthermore, our analyses revealed that combination treatment and, to a smaller degree, quisinostat monotherapy result in significant downregulation of genes involved in the repair of DNA DSB, cell division and chromatin organization (Supplemental Figure 5E). Our results thereby establish that quisinostat, especially when combined with radiation treatment, induces strong transcriptional changes in tumors that promote cellular differentiation, cell cycle arrest and cell death (Supplemental Figure 5).

Interestingly, in end-stage tumors we found that combination treatment induces striking upregulation of multiple genes associated with neuronal development, growth, function, and identity (e.g. axon guidance, axonal fasciculation, neuronal action potential, regulation of dopamine secretion, telencephalon development, GABA signaling pathway; Figure 6D). Similar changes were more modestly induced by quisinostat monotherapy, but not by radiation therapy (Figure 6D). These results strongly suggest that quisinostat treatment causes glioma cells to differentiate and adopt a neuronal-like cell fate, and that these transcriptional changes are significantly amplified when the drug is combined with radiation therapy (Figure 6D). Moreover, GO analysis of downregulated genes demonstrate that out of all treatment groups, combination treatment significantly downregulates genes associated with transcription, DNA replication, the cell cycle, chromatin remodeling and DNA repair (Figure 6E). Expression of the top upregulated genes was validated by RT-qPCR confirming that combination and quisinostat-treated tumors upregulated genes that are normally in expressed neurons (*GABRA2*, *CECR2*, *SCN3A*, *NGFR*, *KCNQ3* and *BDNF*). Taken together, these analyses suggest that combination treatment reduces cell proliferation and induces GBM cells to differentiate towards a neuronal lineage (Figures 6D-F, Figure 7).

**Figure 7.**
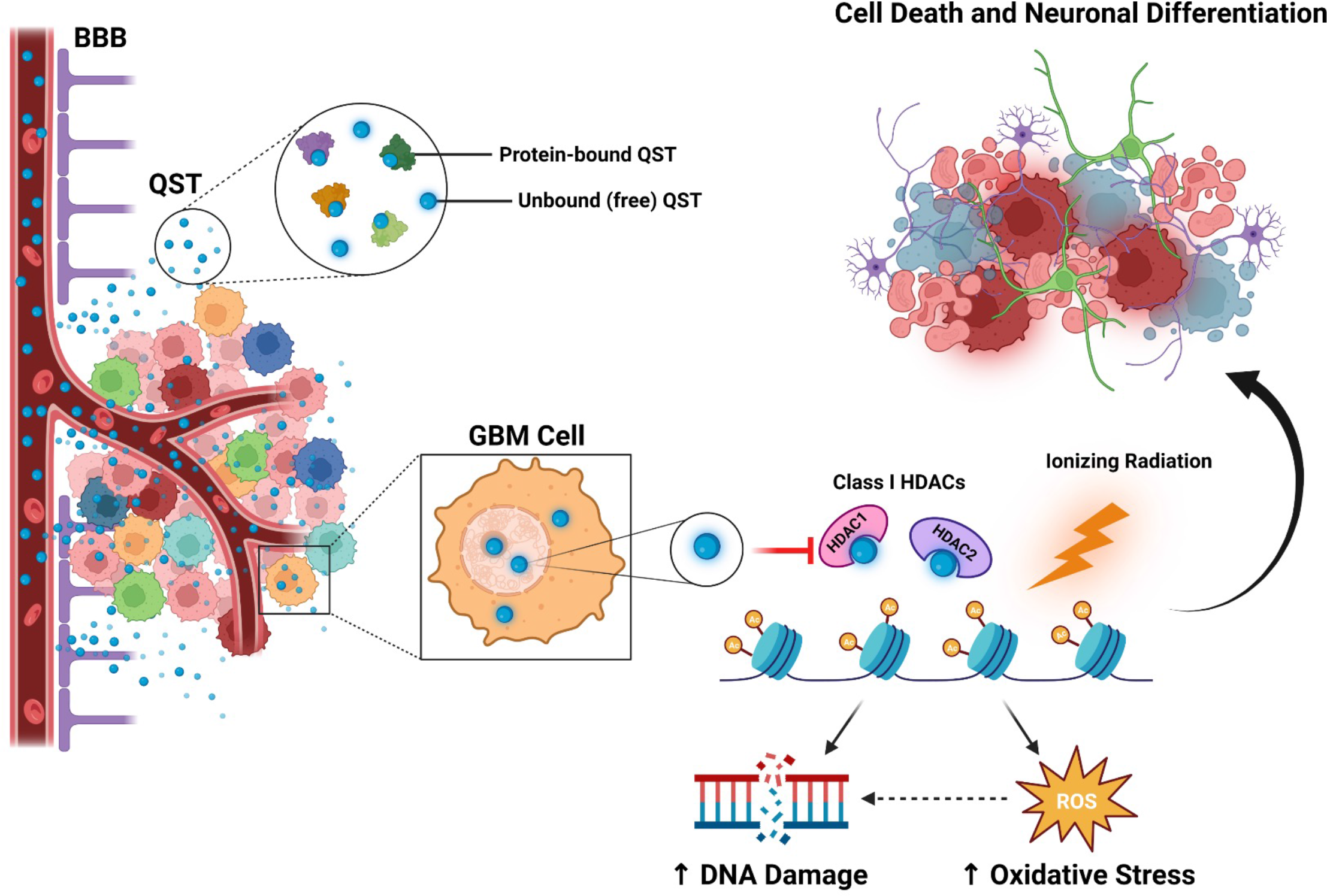
Quisinostat is a brain-penetrant HDACi that sensitizes GBM cells to radiation treatment. Summary of the major findings in this study. Through a careful PK/PD-guided approach, we determined that quisinostat (QST) can cross the blood brain barrier (BBB) and exert its intended PD effect (increase in histone acetylation) in normal brain tissue as well as in GBM cells. As a second-generation HDACi, QST has high sub-nanomolar isoform selectivity for HDAC1 and HDAC2, which are class I HDAC isoforms that are primarily responsible for mediating histone deacetylation. The concentrations of free (non-protein-bound) QST inhibit the function of HDAC1 and HDAC2, resulting in widespread histone hyperacetylation in GBM cells. QST treatment alone in GBM cells also results in increased levels of DNA damage and oxidative stress. When QST is combined with ionizing radiation treatment, genes involved in DNA damage repair pathways and cell division are downregulated, while genes that regulate neuronal development, differentiation and function are significantly upregulated. These findings suggest that combination therapy of QST and radiation provides therapeutic benefit through decreased cell proliferation, dampening of pathways involved in the DNA damage repair response, and increased cell differentiation towards a neuronal-like cell fate.

## DISCUSSION

Systemic inhibition of HDACs is one of the branches of epigenetic therapy that has been investigated in clinical trials for GBM but has so far yielded vastly disappointing results (2, 35, 36). Vorinostat, romidepsin, and panobinostat are the only pan-HDACi that have been tested clinically for GBM as either monotherapies or combination therapies with radiation and/or temozolomide or bevacizumab. All three were found to be ineffective and did not provide a survival benefit when compared to historical control data from previous phase II clinical trials (37-39). The clinical failure of HDACi for the treatment of GBM can be attributed to the fact that most of these drugs are unable to cross the blood-brain barrier at significant concentrations, have high toxicity profiles and thereby small therapeutic windows (2, 40, 41). Notably, these HDACi advanced to clinical trials without PK and target-modulation analyses in the CNS of preclinical models of aggressive gliomas (42-45) – all of which are valuable data necessary for effective clinical translation (42-44).

Several second-generation HDACi have been developed with higher isoform selectivity with the aim of decreasing toxicity and increasing specificity that warrant preclinical investigation for the treatment of GBM (8). Considering that it is widely established that individual HDAC enzymes harbor non-redundant, isoform-specific roles in different kinds of cancers, it is hypothesized that HDACi with greater isoform-selectivity may possess a higher therapeutic index and cause fewer adverse effects (46). As we recently discovered that HDAC1 promotes the tumorigenic properties and survival of glioma stem cells, we questioned whether a brain-penetrant HDACi with higher affinity to HDAC1 would be effective in slowing tumor growth *in vivo*. To this end, we investigated the translational potential of quisinostat – an HDACi that is more selective against class I HDACs and exhibits marked potency towards HDAC1 and HDAC2 – in preclinical models of GBM. We performed a comprehensive PK-PD correlation analysis for quisinostat in preclinical *in vitro* and *in vivo* models of GBM, a study that is first in its kind for any HDACi studied in the field of neuro-oncology.

Here, we demonstrate that quisinostat is highly cytotoxic to human GSC cultures and functions as a potent radiosensitizer in an orthotopic PDX model of GBM. Importantly, we performed brain and tumor tissue-specific PK analyses of total and unbound quisinostat concentrations and demonstrate that it is a brain-penetrant molecule. Our findings are significant considering the importance of HDACi in cancer therapy in general and the controversy surrounding the efficacy of quisinostat for the treatment of malignant brain tumors. One study in a transgenic murine model of sonic hedgehog-driven medulloblastoma reported that quisinostat treatment had a statistically significant but modest survival benefit, while studies in a syngeneic model of GBM and a PDX model of diffuse intrinsic pontine glioma (DIPG) reported no therapeutic benefit (14-16). It was previously suggested that the failure of quisinostat to prolong survival *in vivo* could be attributed to its inability to cross the blood-brain-barrier, but these claims were made in the absence of direct drug level measurements and lack of evidence of on-target engagement in the brain (14, 15). To this end, in our study we utilized an elaborate PK-PD paired analysis approach to establish the brain-penetrant properties of quisinostat and guide interpretation of the translational potential of this HDACi in GBM. We show that when dosed at 10 mg/kg via IP injection, unbound levels of quisinostat can be detected in normal brain tissue and that this concentration is sufficient to induce significant histone H3 hyperacetylation. These results provide experimental evidence that quisinostat can cross a structurally intact blood-brain barrier and induce its intended PD effect in the CNS in contrast to previous conclusions based on survival benefit alone.

We show that while quisinostat significantly slows the growth of intracranial GBM, monotherapy only results in a very modest survival benefit. However, considering that numerous HDACi have previously been shown to display radiosensitizing properties in other malignancies (47-51), we questioned whether quisinostat could enhance radiation-induced cell death in GBM. Indeed, we found that low nanomolar doses of quisinostat robustly synergized with ionizing radiation across multiple glioma cell lines. Importantly, we demonstrate that combinatorial therapy of quisinostat and fractionated radiation resulted in a significant extension of median survival compared to untreated and monotherapy regimens. To the best of our knowledge, this is the first report demonstrating that quisinostat can act as a potent radiosensitizer *in vitro* and *in vivo* in any preclinical cancer model. Hence, our results have important implications for the management of other malignancies – such as prostate, colon, lung, and esophageal cancer – wherein class I HDACs are frequently overexpressed and where radiation is commonly used as a treatment modality (52, 53). Furthermore, our results strongly suggest that quisinostat should be re-evaluated as a potential radiosensitizer in preclinical models of DIPG, given that radiation is currently the only treatment option available for children diagnosed with this aggressive and fatal glioma (54). Our results also strongly suggest that combinatorial treatment of quisinostat and ionizing radiation not only represses genes involved in cellular proliferation, but also induces neuronal differentiation *in vivo*. In line with our observations, a recent study in preclinical models of DIPG also demonstrated that a bifunctional LSD1/HDAC inhibitor (Corin) induces the expression of genes related to neuronal differentiation (55).

We found that quisinostat treatment resulted in elevated levels of DNA double-stranded breaks (DSBs) in glioma stem cells, both *in vitro* and *in vivo*. It is well established that other hydroxamic acid-based HDACi (TSA, vorinostat, panobinostat, belinostat) can induce DNA damage and negatively regulate the different pathways of DNA damage response (DDR) (52, 56-59). While the precise mechanisms through which HDACi directly induce DNA damage and synergize with radiation remain unclear, several hypotheses have been proposed to explain the basis of these phenomena. With respect to the direct induction of DNA damage, previous studies have shown that HDACi can lead to the accumulation of ROS which can result in oxidized DNA base lesions (60-62). If left unrepaired, the oxidative stress-induced single-strand DNA breaks can be converted to DSB during DNA replication (63, 64). Numerous studies have shown that treatment with HDACi across various cancer cell lines also results in the transcriptional downregulation of genes involved in homologous recombination and non-homologous end-joining (*Ku70*, *Ku86*, *DNA-PKcs*, *RAD51*, *BRCA1* and *BRCA2*), which are critical for DSB repair (26, 52, 61). In line with these previous findings, our RNA-seq analysis of tumors receiving both quisinostat and radiation revealed that this combinatorial regimen resulted in the downregulation of genes involved in double-strand DNA repair, homologous recombination, and increased response to oxidative stress. These data suggest that HDACi-induced ROS generation and dampening of the DDR may contribute to the accumulation of DSBs in GBM cells.

Another proposed mechanism for DNA damage is through histone hyperacetylation from HDACi treatment resulting in drastic structural changes in chromatin, exposing large portions of DNA to radiation or other chemotherapeutic agents (65, 66). Hence, it is hypothesized that combination treatment of HDACi and radiation synergize by inducing excessive DNA damage and subsequent apoptosis (28, 67). It is also worth noting that both HDAC1 and HDAC2 have been shown to harbor important roles in the DDR by promoting DSB repair (29, 68). One seminal study demonstrated that HDAC1 and HDAC2 localize to DSB sites and induce local chromatin condensation through deacetylation of histone marks H3K56 and H4K16, repressing transcription and preventing transcription from interfering with DNA repair processes (29). Depletion of both HDAC1 and HDAC2 rendered cancer cells hypersensitive to ionizing radiation and resulted in diminished DSB repair capacity, particularly by non-homologous end-joining (29). Indeed, we found that quisinostat treatment alone led to a gradual significant accumulation of DNA DSBs, and that these effects were further exacerbated when combined with radiation. Hence, we speculate that the radio-sensitizing effects of quisinostat in GSCs may be partly driven through potent inhibition of HDAC1 and HDAC2, against which quisinostat exhibits the highest isoform selectivity (IC_50_: 0.1 nM and 0.3 nM, respectively) (8).

An important limitation inherent in the use of our PDX models is that they preclude an understanding of how an intact immune system may affect the response to quisinostat when combined with radiation treatment. Brain irradiation is known to induce widespread and chronic neuroinflammation which can compromise blood-brain-barrier integrity, cognition, and cell survival (69-71). Hence, it will be necessary to study the radiosensitizing effects of quisinostat in the context of a brain harboring a fully functioning immune system either in syngeneic mouse models of GBM or patient-derived gliomas stem cells transplanted into humanized mouse models.

To date, no hydroxamic acid-based HDACi has been shown to harbor radiosensitizing properties in preclinical models of GBM. However, it remains unclear whether these inhibitors failed to provide any therapeutic benefit due to inadequate brain penetration and/or insufficient on-target modulation. Our study therefore emphasizes the importance of implementing a PK-PD guided approach when evaluating or developing new drugs for GBM. While PK analyses are now commonly performed in preclinical trials for a variety of different brain tumors, these studies typically only measure the total brain-to-plasma concentration (Kp) ratio as a measure of drug-brain penetration (72-74). However, the value of this ratio is rather limited and may lead to erroneous conclusions as it does not consider the protein- or lipid unbound fraction of drug in the plasma and the brain (75). To this end, we employed an equilibrium dialysis method combined with LC-MS/MS analysis to measure the unbound, or “free”, brain-to-plasma concentration (Kp, uu) which represents the pharmacologically active fraction of a drug. Hence, an accurate measurement of both total and unbound drug concentrations in plasma and brain tissues is necessary to establish accurate PK-PD correlations and to understand whether therapeutically relevant concentrations of the drug are present in a brain tumor. A crucial finding in our study was the identification that quisinostat is highly unstable in mouse plasma, with a short half-life of approximately 1 hour. Conversely, quisinostat was highly stable in human plasma – revealing that quisinostat exhibits distinct stability profiles across different species. Importantly, a similar trend was also observed in the mouse brain, wherein quisinostat was found to be much less stable compared to the human brain. A recent study reported that hydroxamic acids such as quisinostat are common substrates of a family of esterases (carboxylesterases) that are abundantly present in rodent plasma but absent in human plasma (33, 76). Indeed, we confirmed that the addition of BNPP, a specific inhibitor of carboxylesterases, stabilized quisinostat in mouse plasma. Our results therefore highlight a large discrepancy in the metabolic stability between rodent and human species. This is an important consideration for translating preclinical studies to the clinic as such differences may hinder the development of promising drug candidates at the preclinical level. Our data and method development will thereby be a valuable resource and note of caution for future preclinical studies employing quisinostat or similar drugs with species-specific stability, as the drug becomes undetectable in mouse plasma over two hours post-dosing if no precautions are taken during sample preparation.

Quisinostat has been tested in phase I and II clinical trials for lung, ovarian and breast cancer, and was well-tolerated at a maximum-tolerated dose of 12 mg given three times weekly. It should be noted that quisinostat harbors superior clinical tolerability compared to panobinostat, another HDACi currently in trials for aggressive gliomas. Data from phase I trials found that while hematologic toxicities (grade 1 and 2) were rare in patients treated with quisinostat (< 5%), such toxicities were far more common and severe (grade 3 and 4) during treatment with panobinostat (5, 6, 77, 78). While the tolerability of quisinostat in combination with other agents in humans remains unexplored, our preclinical data support that combinatorial treatment with fractionated radiation is well tolerated in mice. Considering quisinostat is an HDACi that has passed phase I clinical trials in several cancers and was found to be well-tolerated in humans, it is a promising candidate for phase 0 trial testing in GBM patients (77). This approach would enable the characterization of the PK-PD relationship of quisinostat in humans and fast-track the development of quisinostat as an adjuvant to radiation therapy in phase II trials for GBM.

The identification of drugs that can enhance the effects of radiation treatment is an intense area of research within the field of neuro-oncology, especially for patients with MGMT unmethylated tumors. Nevertheless, while the use of radiosensitizers represents a promising strategy in GBM, the development of these novel agents has been underwhelming (79). Here, we provide the first report that quisinostat is a brain-penetrant HDACi with potent radiosensitizing properties in preclinical models of GBM. Future investigation is required to dissect the molecular consequences of quisinostat treatment and its synergistic relationship with radiation-induced DNA damage in GSCs. Overall, our results provide a rationale for developing quisinostat as a potential combination therapy with radiation for the treatment of GBM.

## METHODS

### Primary Cell Culture

Patient-derived glioma stem cell lines (GSCs; GB187, GB239, GB282, GB7, GB82 and GB126) were established from resected primary GBM tumor tissue at the Barrow Neurological Institute. Cell lines generated at the Barrow Neurological Institute were genetically profiled for mutations and copy number aberrations using the IvySeq custom gene panel established at the pharmacodynamics core at the Ivy Brain Tumor Center. BT145 GSCs were obtained from Dr. Keith Ligon’s laboratory at the Dana-Farber Cancer Institute. All human GSCs were cultured as described previously. U87-MG cells (HTB-14) were purchased from the American Type Culture Collection (ATCC). GSCs were cultured as spheres on non-tissue culture-treated 10cm plates or as adherent cultures on laminin on tissue culture-treated 10 cm plates (ThermoFisher Scientific). GSCs were grown in DMEM/F12 media, supplemented with B27, N2 (Invitrogen, ThermoFisher Scientific) 1% penicillin-streptomycin in the presence of 20 ng/ml epidermal growth factor (EGF) and basic fibroblast growth factor (bFGF) (MilliporeSigma). U87-MG cells were grown in DMEM (Corning) supplemented with 10% BCS (Invitrogen, Gibco, Thermo Fisher) and 1% penicillin-streptomycin (Gibco) on tissue culture-treated 10 cm plates according to manufacturer recommendations.

### Cell Viability Assays After Quisinostat Treatment

GSCs were seeded in laminin-coated tissue culture-treated 96-well plates (clear bottom, white plate; Corning) at a density of 1,000-5,000 cells per well (cell line dependent) in GSC media. U87-MG were seeded using their normal growth conditions without laminin (10% BCS in DMEM). All cells were incubated at 37°C and allowed to adhere overnight. The next day, cells were treated with incremental concentrations of quisinostat (Selleckchem; 0, 10, 25, 50, 100, 250, 500, 1000 nM) diluted in media. Cells treated without quisinostat were treated with DMSO diluted in media. Following treatment with quisinostat, cells were grown for 3-5 days (cell-line dependent) at which point cell viability was measured and quantified. All cell viability measurements were performed using the *CellTiter-Glo*® Luminescent Cell Viability Assay (Promega) following the manufacturer’s instructions. All cell viability results represent the mean of at least 2 biological replicates, each containing three technical replicates.

### Western Blotting

Cellular protein from cultured cells were homogenized in RIPA lysis buffer containing protease and phosphatase inhibitors (ThermoFisher Scientific), rotated at 4 °C for 20 minutes and then centrifuged at 15,000 rpm for 10 minutes at 4 °C. Protein concentration from whole-cell extracts were determined using the Bradford Protein Assay (ThermoFisher Scientific). Equal amounts of protein (10-40 μg/lane) were loaded onto a 10% or 12.5% SDS-PAGE gels and transferred to a polyvinylidene fluoride membrane (PVDF; Millipore-Sigma).

Cellular protein from frozen tissue of non-tumor bearing mouse brains, tumor-bearing mouse brains and flank tumors were homogenized in a pre-chilled glass tissue grinder (VWR) with RIPA lysis buffer containing protease and phosphatase inhibitors (ThermoFisher Scientific). 500uL of RIPA buffer was used for 10 mg of tissue. Once homogenized, the tissue lysates were kept on ice for 30 minutes and vortexed every 10 minutes. The samples were then centrifuged at 15,000 rpm for 10 minutes at 4 °C to collect the protein lysates.

Membranes were blocked with 5% non-fat milk for 1 hour at room temperature and incubated overnight with primary antibody at 4 °C; Primary antibodies used in this study were rabbit anti-p21 (1:500; Abcam, ab109520), rabbit anti-gamma H2AX (phospho Ser139; 1:1000, Abcam, ab11174), rabbit anti-H3K27ac (2 μg/mL, Abcam, ab4729), rabbit anti-H3K9/14ac (1:1000, Cell Signaling Technologies, 9677), mouse anti-GAPDH (1:1000, Cell Signaling Technologies, 97166) and mouse anti-β-actin (1:1000, Bio-Rad, MCA5775GA). Membranes were probed with fluorophore-conjugated anti-mouse or anti-rabbit secondary antibodies (1:10,000; ThermoFisher Scientific). Western blots were developed using the LI-COR Odyssey CLx imaging system (LI-COR Inc.) and quantitated using the Image Studio Lite software. All Western blots are representative images from a minimum of three biological replicates.

### Immunocytochemistry

Cells were grown as adherent cultures on laminin-coated glass coverslips (Thermo Fisher Scientific) in GSC media. 24 hours after plating the cells were treated with quisinostat or DMSO diluted in GSC media. 72 hours post-treatment, cells were and fixed with 4% paraformaldehyde (PFA) for 13 minutes at room temperature. Cells were washed with PBS and subsequently permeabilized and blocked with 5% normal goat serum (Sigma Aldrich) and 0.2% Triton X-100 in PBS (blocking solution) for 30 minutes at room temperature. The cells were incubated with primary antibodies overnight at 4°C in blocking solution. Primary antibodies used in this study included rabbit anti-Ki67 (1:1000; Abcam, 15580), rabbit anti-Cleaved Caspase 3 (1:400; Cell Signaling Technologies, 9661), rabbit anti-gamma H2AX (phospho Ser139; 1:1000, Abcam, ab11174), and mouse anti-human Nestin (1:500; Novus Biologicals, 10C2). The following day, the cells were washed with PBS three times, incubated with fluorophore-conjugated secondary antibodies at 1:1,000 dilutions (Alexa Fluor 568 goat anti-mouse, Abcam, ab175473; Alexa Fluor 488 goat anti-rabbit, Abcam, ab150077) for 1 hour at room temperature, and finally washed in PBS three more times. Cells were mounted onto SuperFrost Plus microscope slides using Fluoroshield Mounting Medium containing DAPI (Abcam). Images were acquired using a confocal microscope (Leica Microsystems; TCS SP5) operated with LAS software. The fraction of Ki67- and Cleaved Caspase 3-positive cells were counted from five independent images from each condition. The average and standard deviation were calculated from three biological replicates for all control and Quisinostat-treated experiments.

### Image Acquisition

Analysis of immunostaining of cultured GSCs were performed on confocal stacks (with a step size of 0.5-1.5 μm) acquired with a either a 20x water-immersion objective or a 63x oil-immersion objective on a laser-scanning confocal microscope (Leica Microsystems; TCS SP5) operated with LAS software. All images were processed using the ImageJ software (NIH).

### In vitro Irradiation Studies

For all *in vitro* radio-sensitization experiments involving treatment with ionizing irradiation (IR) using RS 2000 irradiator (Rad Source) or the X-Rad225XL Irradiator (Precision X-Ray), GSCs were plated on laminin-coated tissue culture-treated 96-well plates and incubated at 37°C overnight for 24 hours. The next day the cells were pre-treated with quisinostat or an equivalent volume of DMSO for one hour and then subsequently irradiated with various doses of IR (cell-line dependent). Cell viability was measured as described above using the *CellTiter-Glo*® assay (Promega) 3-5 days after treatment. For experiments involving protein characterization of IR-treated cells preceded by treatment with quisinostat, whole-cell lysates were collected 1, 2, 6 and 24 after irradiation. Radiation was delivered using a RS2000 Series Biological Irradiator (Rad Source Technologies) or the X-Rad225XL Irradiator (Precision X-Ray),

### Flank Tumor Implantation

For flank implantations, the cells were prepared in a 1:1 ratio with 100 uL Matrigel (Corning #356234) and 100 uL of a single cell suspension of U87 (500,000 cells) in a 1 mL syringe fitted with a 26-gauge needle. The mice were anesthetized with isoflurane in a plastic desiccator placed in an externally vented fume hood. The U87-Matrigel cell suspension was then subcutaneously injected into the flank of the mouse on the posterior/lateral aspect of the lower rib cage. The mice were monitored daily and growth of flank tumor area was measured with a digital caliper (ThermoFisher) once a week. Mice were sacrificed once the tumor size grew over 2000 mm^3^ in size.

### Orthotopic Xenograft Studies

7-week old *Foxn1^nu^* nude male mice (The Jackson Laboratory) were used for *in vivo* orthotopic transplantation of luciferized GB126 (male) cells. Nude mice were anesthetized using gaseous isoflurane and immobilized on a Leica stereotaxic instrument (cat# 39477001, Leica Microsystems). Following an incision at the midline, a fine hole was drilled 2.5mm lateral to the bregma. Using a 33-guage needle syringe (700 series, Hamilton), 2 μl of dissociated viable cells (at a density of 50,000 cells/μl) were injected 2 mm deep from the surface of the skull slowly at a constant rate of 1 μl per minute for 2 minutes. The needle was left for 1 additional minute to prevent reflux of the injected cells and was then slowly removed. The incision was closed with surgical staples. All mice were observed daily and were sacrificed upon the onset of severe neurological symptoms and >10% body weight loss. Survival data was plotted and analyzed using GraphPad Prism 8 (GraphPad Software).

### Live Bioluminescence (IVIS) Imaging

2 weeks post-implantation, the mice were examined for tumor growth by monitoring bioluminescence every 7 days using the IVIS Xenogen Spectrum platform. D-Luciferin Potassium Salt (Gold Biotechnology) was dissolved in PBS at a final concentration of 15 mg/mL. All mice were weighed each week and were administered D-Luciferin via an intraperitoneal injection (10μl/g). 15 minutes after the injection, the mice were sedated using gaseous isoflurane (Piramal) and placed inside an IVIS Spectrum In Vivo Imaging System (Perkin Elmer) for bioluminescence imaging. The total flux (photons/second) within the region of interest (ROI) was calculated using the Living Image Software 4.5 (Perkin Elmer).

### Preparation of Quisinostat for In Vivo Use

For *in vivo* preparation, quisinostat was dissolved in 50% PEG-300, 50% sterile water solution for 10 mg/kg dosing. The suspension was then sonicated for 10 minutes to allow the drug to completely dissolve. Finally, the pH or both the drug and vehicle solutions were adjusted to 7.4 prior to intraperitoneal dosing.

### Determination of Optimal Administration Route

*Foxn1^nu^* nude male mice (The Jackson Laboratory) were used to determine the drug administration route that would result in best quisinostat bioavailability. Three cohorts of mice were treated with a single dose of 10 mg/kg quisinostat delivered through either intraperitoneal, subcutaneous, or oral gavage routes (3 mice per cohort). Following administration of the single dose, approximately 30 µL of blood was drawn from the tip of the tails at the following timepoints: 0.5, 1, 2, 4, 6, 8 and 24 hours. The collected blood was immediately centrifuged at 3,000 rpm for 10 minutes at 4°C to separate the plasma, which was subsequently flash frozen. At the 24-hour timepoint, following the last blood sample collection, the mice were sacrificed and the whole brains from each mouse were dissected and flash-frozen for subsequent analysis.

### In Vivo Irradiation Studies

Intracranial or flank tumor-bearing nude mice were sedated with gaseous isoflurane prior to irradiation. On the first week of treatment, 2 hours after treatment with quisinostat or vehicle the mice were treated with 2 Gy of ionizing radiation on MWF for a total 6 Gy (3 doses). Ionizing radiation was administered with the RS2000 Series Biological Research Irradiator (Rad Source Technologies).

### Treatment of Flank-Implanted mice with quisinostat and/or Radiation

Mice with implanted flank tumors were allowed to grow until the tumor size reached 100 mm^3^ volume. Mice were randomized into groups before treatment and underwent treatment on MWF for the entire duration of the experiment until the tumor volumes exceeded 2000 mm^3^.

Treatment groups included vehicle (50% PEG-300), quisinostat alone (10 mg/kg), flank IR treatment with vehicle (2 Gy), and flank IR treatment (2 Gy) with quisinostat (10 mg/kg). For mice receiving IR treatment with or without quisinostat, mice were treated with 2 Gy on MWF for a total 6 Gy on the first week of treatment. Quisinostat was administered to mice through intraperitoneal injections two hours prior to flank tumor radiation treatment. Upon completion of the radiation regimen, IR-treated mice subsequently received quisinostat or vehicle alone for the rest of the experiment. Tumor growth and treatment response was monitored by manually measuring the tumor area once per week starting at 13 days post-implantation. Upon reaching the 2000 mm^3^ tumor volume threshold, mice were sacrificed, and plasma and tissue samples were harvested for PD and PK analyses two hours after treatment with a final dose of quisinostat (10 mg/kg).

### Treatment of Intracranially-implanted mice with Quisinostat and/or Radiation

Mice with implanted tumors were allowed to grow until the tumor bioluminescence score reached 10^8^ radiance (p/s/cm3/sr). Mice were randomized into groups before treatment. For survival studies, mice underwent treatment on MWF for the entire duration of the experiment until moribund. For mice receiving IR treatment with or without quisinostat, mice were treated with 2 Gy on MWF for a total of 6 Gy on the first week of treatment. Treatment groups included vehicle (50% PEG-3000), quisinostat (10mg/kg) alone, 6 Gy whole brain IR treatment with vehicle and 6 Gy whole brain IR treatment with 10 mg/kg quisinostat. Quisinostat was administered to mice through intraperitoneal injections two hours prior to whole-brain radiation treatment. Upon completion of the radiation regimen, IR-treated mice subsequently received quisinostat or vehicle alone for the rest of the experiment. Tumor growth and treatment response was monitored by IVIS bioluminescence once per week. For survival studies, mice were sacrificed, and samples were collected for PD and PK analyses once moribund two hours after treatment with a final dose of quisinostat (10 mg/kg).

For short-term PK / PD correlation studies, tumor-bearing mice were randomized into groups and underwent a single week of treatment with quisinostat (10 mg/kg) on MWF. For mice receiving IR treatment with or without quisinostat, mice were treated with 2 Gy on MWF for a total of 6 Gy. Quisinostat was administered to mice through intraperitoneal injections two hours prior to whole-brain radiation treatment. On the third and last day of treatment, mice were sacrificed and processed for PD and PK analyses 3 hours after administration of quisinostat or vehicle. For PD analyses, the mice were euthanized with isoflurane and the tumors were resected out of the brain and flash-frozen for subsequent analysis through western blotting and RNA-sequencing. Tissue from the hemisphere contralateral to the tumor was also collected as a normal brain / non-tumor reference sample. For PK analyses, the mice were anesthetized with isoflurane and at least 300 μL of blood was drawn from the right ventricle of the heart. The blood was collected in tubes containing 10 μL of 0.1 M KOH in EDTA to prevent blood coagulation. Blood samples were immediately centrifuged at 3,000 rpm for 10 minutes at 4°C to allow separation of plasma. After collection of the blood, the tumor was dissected out of the brain and flash-frozen. Both plasma and erythrocytes were flash-frozen for subsequent analysis.

### Bioanalytical LC-MS/MS Method

Quisinostat concentrations in specimens were measured using reverse-phase liquid chromatography on the AB SCIEX QTRAP6500+ LC–MS/MS system by operating electrospray in the positive ion mode. For liquid chromatographic separation, gradient elution was performed using a Phenomenex Kinetex F5 100 Å column (100 × 2.1 mm, 2.6 *μ*m). The initial composition of the mobile phase was composed of 70% phase A (0.1% formic acid in water) and 30% phase B (0.1% formic acid in 1:1 acetonitrile:methanol) with a 0.35 ml/min flow rate. Gradient elution was achieved as follows: mobile phase (B) was maintained at 30% from 0 to 0.3 minutes, increased to 95% from 0.3 to 0.8 minutes, maintained at 95% from 0.8 to 2.5 minutes, and lowered to 30% from 2.5 to 2.8 minutes. The total run time was 3.5 minutes. The internal standard used in this study was D8-infigratinib. The retention times for quisinostat and D8-infigratinib were 1.6 and 1.8 minutes, respectively. Mass-to-charge ratio (m/z) transitions were as follows: 395.20 → 144.00 (quisinostat) and 568.08 → 321.00 (D8-infigratinib). LC-MS/MS analysis was performed using Analyst 1.7 Chromatographic Data System (Foster City, CA, USA).

### Calibration standards and quality control samples

Stock solutions of standard (1 mM quisinostat) and IS (250 µM D8-infigratinib) were prepared in acetonitrile. Working solutions for calibration curve standards and quality controls (QC) were prepared by dilutions with a 40% methanol mixture aqueous solution. The IS precipitation solution (10 nM) was prepared from the IS stock solutions by dilution with methanol. Calibration standards and batch qualifying QCs were freshly spiked for every batch. For sample analysis in human and mouse matrices, calibration standards were prepared in bulk by spiking appropriate amounts of working solutions into blank human plasma, used as a surrogate matrix due to the instability of quisinostat in mouse plasma. For sample analysis in neural stem cell (NSC) media and cell lysate, calibration standards were prepared in bulk by spiking appropriate amounts of working solutions into NSC media. QC samples were prepared in bulk by spiking appropriate amounts of working solutions into blank mouse plasma or cell media. Preparation of calibration standards and QC samples was performed at 4°C. Final concentrations range of the calibration standards were 1 – 1000 nM in human plasma or NSC media. Three QC levels, namely low (LQC), medium (MQC), and high (HQC), were used during all sample analyses. The concentrations of QC samples in various matrices were 3 nM (LQC), 22 nM (MQC), and 800 nM (HQC). All stock solutions and working solutions were stored at 4°C.

#### Plasma sample preparation

Frozen plasma samples were thawed at 4°C. An aliquot of 30 µL mouse plasma was transferred into a micro centrifuge tube followed by 30 µL of blank human plasma, and precipitation with 180 µL of IS-containing methanol precipitation solution. The mixture was vortex-mixed for 10 s and centrifuged at 12000 g at 4 °C for 10 min. A 100 µL aliquot of the supernatant was transferred to an autosampler vial and 5 µL was injected into the LC–MS/MS system for analysis. Whenever necessary, appropriate dilutions were made to accommodate the analysis of small volume samples.

#### Brain and brain tumor sample preparation

Normal brain and brain tumor tissue homogenates were prepared by 1:4 (mass/volume) ratio with PBS. Samples were homogenized under 6.00 m/s speed for 40 seconds with three cycles by Bead Ruptor Elite homogenizer (Omni International, USA). Plasma was used as a surrogate matrix for brain/tumor homogenate. Brain homogenate samples from *in vivo* studies were prepared as described for plasma samples. Analyte and IS were extracted by protein precipitation with methanol containing IS. After centrifugation at 12000 rpm for 10 minutes at 4^0^C, 5 µL of supernatant was injected into LC–MS/MS system for analysis.

#### Cell media and lysate sample preparation

Cell media and lysates were thawed at room temperature. An aliquot of 20 µL of cell media or lysate was transferred into a micro centrifuge tube followed by protein precipitation with 60 µL of IS-containing methanol precipitation solution. The mixture was vortex-mixed for 10 s and centrifuged at 12000 g at 4 °C for 10 min. A 50 µL aliquot of the supernatant was transferred to an autosampler vial and 5 µL was injected into the LC–MS/MS system for analysis.

#### Stability Study in Mouse Plasma, Mouse Brain, Human Plasma, Human Brain, and NSC Media

The stability of quisinostat was determined in BALB/c mouse plasma, male nude athymic perfused and non-perfused mouse brain homogenate (1:9 w/v of PBS (pH 7.4)), human brain (obtained from VRL Eurofins, Denver, CO, USA) homogenate (1:4 w/v of PBS (pH 7.4)), pooled human plasma (Innovative Research INC, Novi, MI, USA), and neurobasal media. Quisinostat stock solutions (1 mM) were prepared in acetonitrile, subsequently diluted in a 40% methanol mixture, and added to the matrices to make final concentrations of 100 nM or 10 nM. 50 or 30 µL of either plasma or brain homogenate containing quisinostat were aliquoted into 1.5 mL microcentrifuge tubes (Eppendorf) and were incubated at either 4°C or 37°C for 0, 2, 4, 6, 12, or 24 hours (n = 3 at each time point). The stability of quisinostat in mouse plasma was also tested at 0, 2, 8, 16, or 24 hours (*N* = 3 at each time point) in the presence of esterase inhibitors: 0.25 mM bis(p-nitrophenyl) phosphate (BNPP), 1mM dithiobis(2-nitrobenzoic acid) (DTNB), or 1.25 mM phenylmethylsulfonyl fluoride (PMSF). Both plasma and brain homogenate samples were stored at −80°C until liquid chromatography–tandem mass spectrometry (LC-MS/MS) analysis.

### qRT-PCR

Total RNA was extracted from orthotopic tumors by using the PureLink RNA Mini Kit (Ambion) in accordance with the manufacturer’s instructions. RNA was quantified on a NanoDrop Spectrophotometer (Tecan), and 1 μg of total RNA was used for cDNA synthesis by using the SuperScript VILO kit (Life Technologies). qPCR was performed using inventoried TaqMan assays for respective target genes and housekeeping control genes (18S) on the QuantStudio 6 Flex Real-Time PCR System (Life Technologies). Fold change in gene expression was analyzed using the delta delta Ct method.

### RNA-seq analysis

To analyze RNA-seq data, raw RNA-seq reads were selected for quality and length by removing low-quality reads and adapter sequences using cutadapt. Samples were then aligned to two separate genomes, GRCm39 (Mouse) and GRCh38.p13 (Human) using STAR (80). Samples were then filtered with XenofilteR using Human bam as Graft and Mouse as Host to reduce mouse DNA in samples compared. Count tables were generated using featureCounts in the Subread package (81) and the resulting count tables were analyzed in R using DESeq2 to identify differentially expressed genes. Genes that were up- or down-regulated at least two-fold with an FDR <0.05 were considered differentially expressed for downstream analyses. After identifying differentially expressed genes, Gene Ontology analyses (34) were performed using the Fisher method (cutoff p<0.01) to identify gene categories that were up- and down-regulated by the various treatments.

### Statistical Analysis

Data are presented as the mean ± SEM. If comparing two conditions or cell lines, significance was tested with unpaired two-tailed Student’s t-test. Significance of the differences between conditions or cell lines were tested by the two-way ANOVA with Bonferroni multiple comparison tests using GraphPad Prism 9 (GraphPad software). Survival studies were analyzed using the Kaplan-Meier method with the Mantel-Cox log-rank test (GraphPad software). Statistical significance was defined at * *p* < 0.05, ** *p* < 0.01, *** *p* < 0.001, **** *p* < 0.0001.

### Study Approval

The patient samples used for this research were provided by the Biobank Core Facility at St. Joseph’s Hospital and Medical Center and Barrow Neurological Institute (BNI). The samples were de-identified and conformed to the Biobank Institutional Review Board’s protocol. Animal husbandry was performed in accordance with the guidelines of the St. Joseph’s Hospital and Medical Center and Barrow Neurological Institute under the protocol approved by the Institutional Animal Care and Use Committee.

## ACKNOWLEDGEMENTS

Patient-derived glioma cells were provided by the Biobank Core Facility at St. Joseph’s Hospital and BNI and the Living Tissue Bank at Dana-Farber Cancer Institute. The biobank is funded by the Arizona Biomedical Research Commission and the Barrow Neurological Foundation. We are grateful to Keith Ligon for providing several patient-derived glioma cell lines. We thank the entire Mehta Lab and An-Chi Tien for helpful discussions, technical assistance, and manuscript comments. We thank MOGene for assistance with bioinformatics and analysis of RNA-seq data. The schematics in Figures 4, 5 and 7 were generated on BioRender.com. We thank the Students Supporting Brain Tumor Research (SSBTR) charity for supporting this work. This work was supported by grants to SM from the NIH National Institute of Neurological Disorders and Stroke (NINDS) (R01 NS088648A) and to SM, AT, and NS from the Barrow Neurological Foundation. NS is supported by grants from the NIH/National Cancer Institute (R01 CA175391). SM, AT, and NS are supported by the Ben and Catherine Ivy Foundation. Funding sources were NIH/NINDS, Barrow Neurological Foundation, and the Ben and Catherine Ivy Foundation.

## AUTHOR CONTRIBUTIONS

CLC, AT and SM conceived and designed all experiments. CLC, ELM, JBM and CIW standardized the techniques, performed in vitro and in vivo experiments, and analyzed the data. TM, WK and AT performed all analyses and developed methods for pharmacokinetic studies. SG performed in vitro experiments and assisted with data analysis. ZO assisted with in vivo experiments. WY performed and helped with statistical analyses. TM, ELM, JBM, CIW, ZO, NS and AT edited the manuscript. NS provided patient tissues to establish patient-derived glioma cell lines. SM and AT coordinated and supervised the project. CLC and SM wrote the manuscript.

---

**Supplemental Figure 1. Genomic characterization of cell lines used in this study.** Patient-derived GSCs were sequenced and profiled for genetic mutation and copy number variation aberrations using the IvySeq custom gene panel developed at the Ivy Brain Tumor Center. (P = primary GBM; R = recurrent GBM).

**Supplemental Figure 2. Temporal dynamics of intracellular uptake of quisinostat in BT145.** (A) Levels of intracellular quisinostat with continuous exposure to drug (75 nM) in BT145 over the course of 24 hours. Blue line indicates measured intracellular levels of quisinostat at 2, 6, 10, and 24 hours after treatment. Orange line with triangles indicates levels of quisinostat present in the cell media at each collected timepoint. Orange line with circles denotes the baseline levels of drug present when it was spiked into the cell media for each timepoint (t = 0 hr). (B) Levels of intracellular quisinostat in BT145 cells that were treated with 75 nM quisinostat and underwent drug washout 2 hours after initial treatment. Blue line indicates measured intracellular levels of quisinostat at 2, 6, 10, and 24 hours after drug washout. Orange line with triangles indicates levels of quisinostat present in the cell media at each collected timepoint. Orange line with circles denotes the baseline levels of drug present when it was spiked into the cell media for each timepoint (t = 0 hr). QST = quisinostat. The data are compiled from at least three independent experiments. Error bars indicate SEM.

**Supplemental Figure 3. Quisinostat synergizes with ionizing radiation in vitro.** (A-B) Matrices illustrating the BLISS and Loewe synergy scores when combining quisinostat (0-1000 nM) with increasing doses of radiation in (A) BT145 and (B) GB126. Related to Figure 2G-H.

**Supplemental Figure 4. Stability of quisinostat in mouse plasma is improved upon addition of esterase inhibitors.** (A) Stability of quisinostat (100 nM) in mouse plasma and mouse brain homogenate when sample preparation is performed at 4°C. (B) Stability of quisinostat (500 nM) in mouse plasma spiked with three different esterase inhibitors (BNPP, PMFS, DTNB) over the course of 24 hours. (C) Stability of quisinostat (10 nM and 100 nM) in neurobasal cell culture medium over the course of 24 hours.

**Supplemental Figure 5. RNA-seq analysis of short-term (acute) quisinostat treatment in vivo.** (A-B) Venn diagrams showing the overlap in genes either upregulated (A) or downregulated (B) in response to quisinostat (QST) monotherapy, radiation treatment (IR) or combination treatment (QST + IR). Mice received only three doses in total and were sacrificed 3 hours post-dosing. Gene numbers in each section are shown in parentheses. (C) Volcano plots showing the –log10 (p value) and log2 fold change for transcripts detected by RNA-seq analysis of acutely-treated tumors treated with QST (left), IR (center), or QST + IR (right). Significantly up- and downregulated genes (FDR < 0.05, 2-fold) are marked in red and blue, respectively. (D- E) Gene ontology analysis of genes upregulated (D) or downregulated (E) in GB126 tumors due to QST, IR or QST + IR treatment.

