## Supplemental Figures 1-5 for "Quisinostat is a brain-penetrant radiosensitizer in glioblastoma"

### Supplemental Figure 1

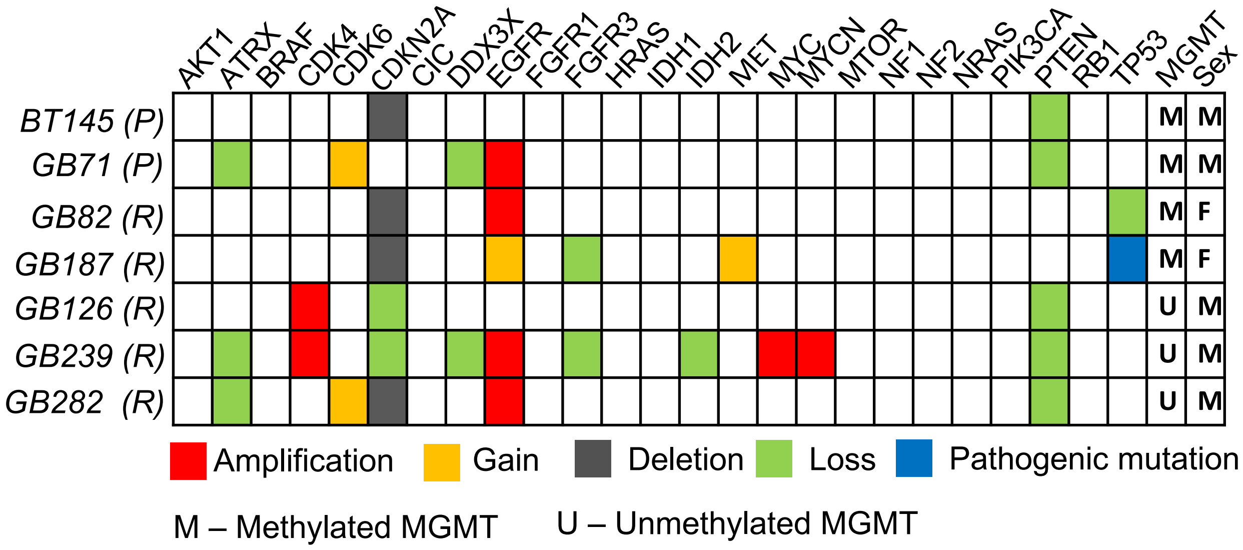


**Supplemental Figure 1. Genomic characterization of cell lines used in this study.** Patient-derived GSCs were sequenced and profiled for genetic mutation and copy number variation aberrations using the IvySeq custom gene panel developed at the Ivy Brain Tumor Center. (P = primary GBM; R = recurrent GBM).

**Supplemental Figure 2**

A.

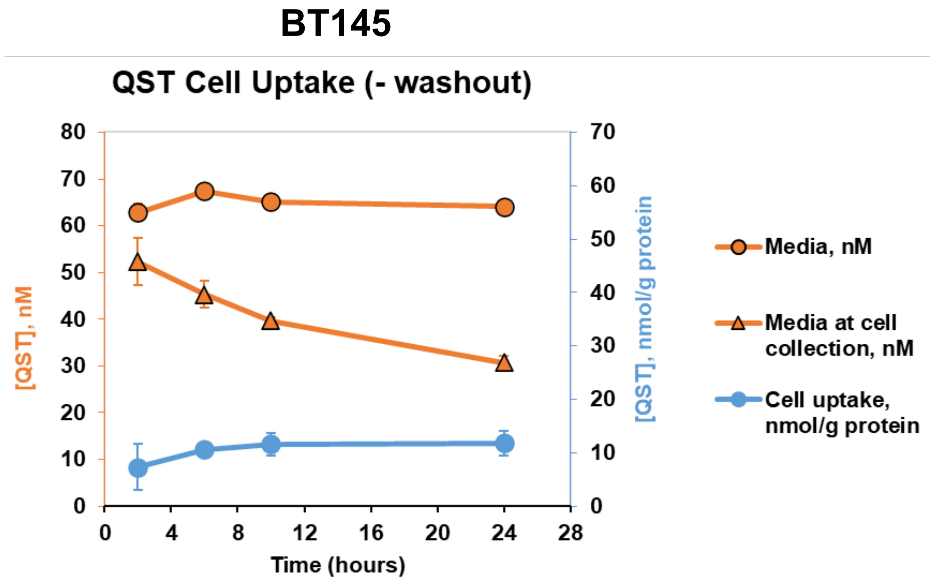


B.

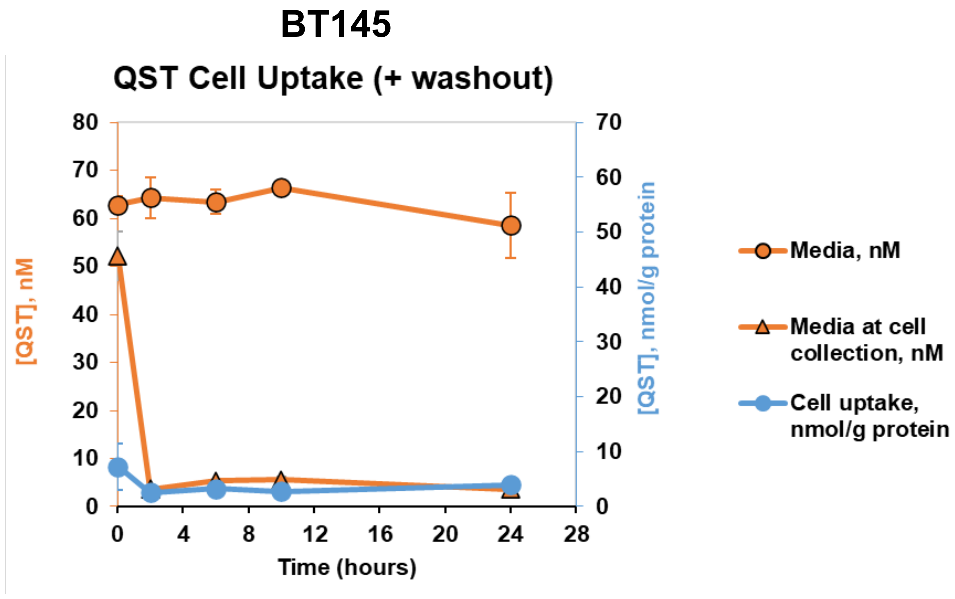


**Supplemental Figure 2. Temporal dynamics of intracellular uptake of quisinostat in BT145.** (A) Levels of intracellular quisinostat with continuous exposure to drug (75 nM) in BT145 over the course of 24 hours. Blue line indicates measured intracellular levels of quisinostat at 2, 6, 10, and 24 hours after treatment. Orange line with triangles indicates levels of quisinostat present in the cell media at each collected timepoint. Orange line with circles denotes the baseline levels of drug present when it was spiked into the cell media for each timepoint (t = 0 hr). (B) Levels of intracellular quisinostat in BT145 cells that were treated with 75 nM quisinostat and underwent drug washout 2 hours after initial treatment. Blue line indicates measured intracellular levels of quisinostat at 2, 6, 10, and 24 hours after drug washout. Orange line with triangles indicates levels of quisinostat present in the cell media at each collected timepoint. Orange line with circles denotes the baseline levels of drug present when it was spiked into the cell media for each timepoint (t = 0 hr). QST = quisinostat. The data are compiled from at least three independent experiments. Error bars indicate SEM.

### Supplemental Figure 3

## A.

### BT145

BLISS LOEWE

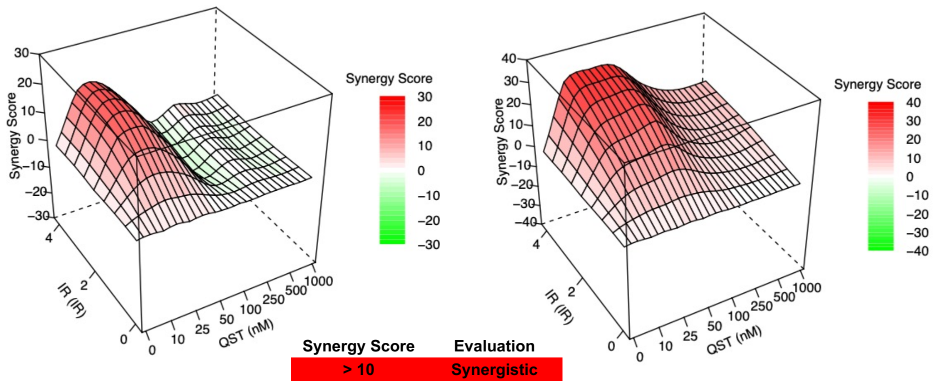


| Synergy Score | Evaluation |
|---|---|
| > 10 | Synergistic |
| -10 to 10 | Additive |
| < -10 | Antagonistic |

## B.

### GB126

BLISS LOEWE

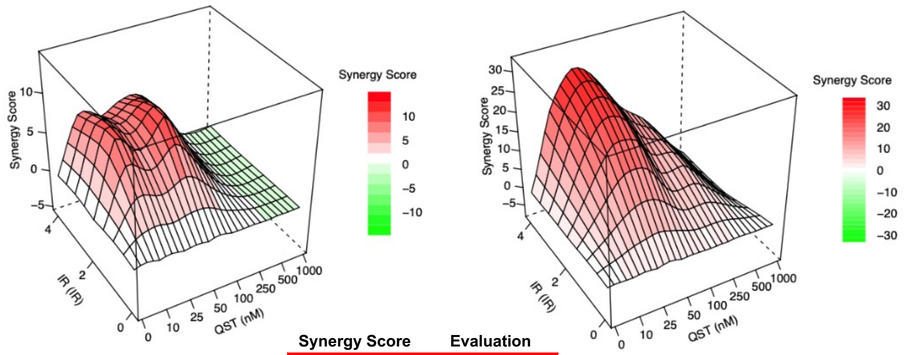


| Synergy Score | Evaluation |
|---|---|
| > 10 | Synergistic |
| -10 to 10 | Additive |
| < -10 | Antagonistic |

**Supplemental Figure 3. Quisinostat synergizes with ionizing radiation *in vitro*.** (A-B) Matrices illustrating the BLISS and Loewe synergy scores when combining quisinostat (0-1000 nM) with increasing doses of radiation in (A) BT145 and (B) GB126. Related to Figure 2G-H.

### Supplemental Figure 4

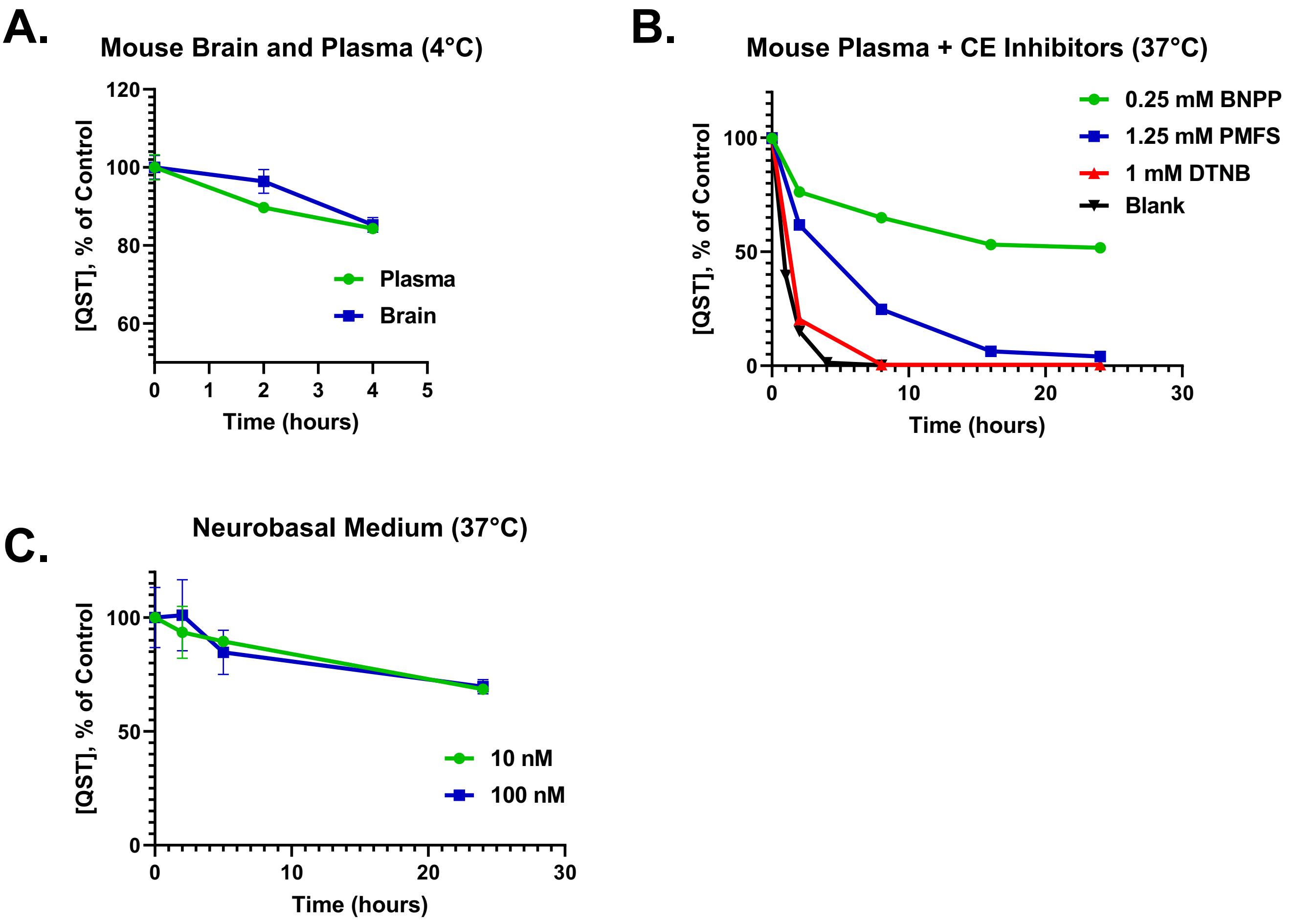


**Supplemental Figure 4. Stability of quisinostat in mouse plasma is improved upon addition of esterase inhibitors.** (A) Stability of quisinostat (100 nM) in mouse plasma and mouse brain homogenate when sample preparation is performed at 4°C. (B) Stability of quisinostat (500 nM) in mouse plasma spiked with three different estserase inhibitors (BNPP, PMFS, DTNB) over the course of 24 hours. (C) Stability of quisinostat (10 nM and 100 nM) in neurobasal cell culture medium over the course of 24 hours.

**Supplemental Figure 5**

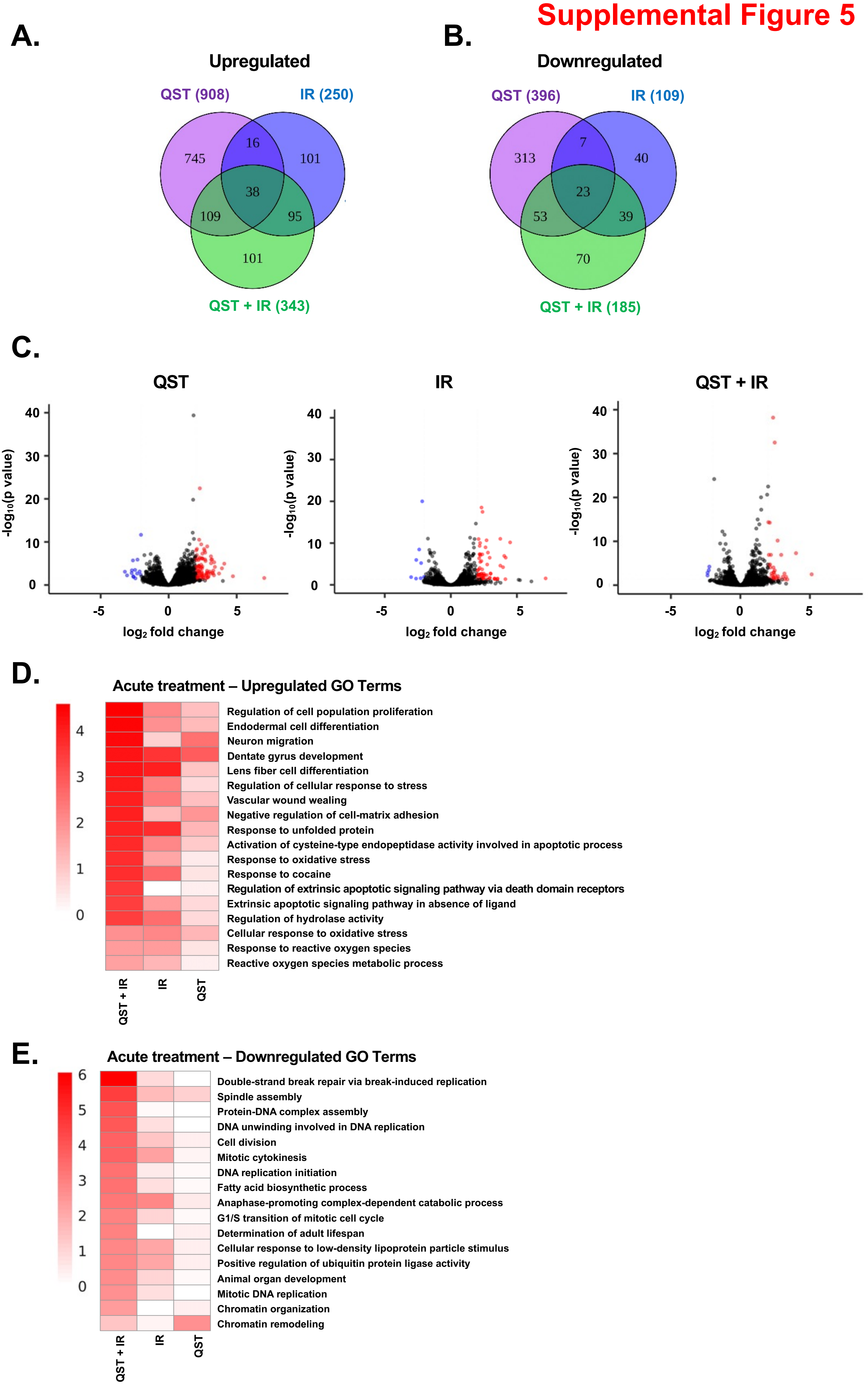


**Supplemental Figure 5. RNA-seq analysis of short-term (acute) quisinostat treatment *in vivo*.** (A-B) Venn diagrams showing the overlap in genes either upregulated (A) or downregulated (B) in response to quisinostat (QST) monotherapy, radiation treatment (IR) or combination treatment (QST + IR). Mice received only three doses in total and were sacrificed 3 hours post-dosing. Gene numbers in each section are shown in parentheses. (C) Volcano plots showing the –log10 (p value) and log2 fold change for transcripts detected by RNA-seq analysis of acutely-treated tumors treated with QST (left), IR (center), or QST + IR (right). Significantly up- and downregulated genes (FDR < 0.05, 2-fold) are marked in red and blue, respectively. (D-E) Gene ontology analysis of genes upregulated (D) or downregulated (E) in GB126 tumors due to QST, IR or QST + IR treatment.
